# CHD4 and NKX2.2 Cooperate to Regulate Beta Cell Function by Repressing Non-Beta Cell Gene Programs

**DOI:** 10.1101/2025.06.16.659956

**Authors:** Dylan Sarbaugh, Thais Gaia Oliveira, Michelle A. Guney, McKenna R. Casey, Victoria M. Hoelscher, Christopher J. Hill, Cole R. Michel, Kristen L. Wells, Richard K.P. Benninger, Lori Sussel

**Affiliations:** Barbara Davis Center for Diabetes, University of Colorado Anschutz Medical Campus, Aurora, CO 80045, USA; School of Pharmacy, Mass Spectrometry Facility, University of Colorado Anschutz Medical Campus, Aurora, CO 80045, USA

## Abstract

NKX2.2 is a transcription factor that regulates pancreatic islet beta (β) cell identity and function; however, cofactor proteins that modulate the functional activity of NKX2.2 in β cells are relatively unexplored. An unbiased proteomics screen identified chromodomain helicase DNA-binding protein 4 (CHD4) as an NKX2.2 interacting partner. CHD4 is a nucleosome remodeler that directs the appropriate differentiation, maturation and function of many cell types. To characterize the roles of CHD4 in β cells, we generated *Chd4* βKO mice. Deletion of *Chd4* substantially impaired the function of β cells. The *Chd4* βKO mice became diabetic due to the disruption of islet integrity, calcium signaling and downregulation of essential β cell regulatory genes. We also discovered CHD4 is required to bind at and repress non-beta cell genes, including *Kcnj5,* the gene that encodes the GIRK4 potassium channel in β cells. Aberrant upregulation of GIRK4 causes impaired glucose-stimulated insulin secretion. These studies demonstrate that CHD4 is an essential transcriptional cofactor of NKX2.2 that is required for the proper maturation and function of pancreatic β cells.

**Article Highlights:**

- NKX2.2 interacts with the Nucleosome Remodeling and Deacetylase (NuRD) complex through its interaction with CHD4.
- Deletion of CHD4 from developing pancreatic β cells in mice causes diabetes due to a loss of islet integrity, disrupted calcium signaling and impaired insulin secretion.
- Beta cells lacking CHD4 inappropriately upregulate the GIRK4 potassium channel; inhibition of GIRK4 rescues the insulin secretion defect.

## Introduction

Blood glucose homeostasis is maintained by the pancreatic islets of Langerhans and when pancreatic β cells become dysfunctional or destroyed, diabetes mellitus ensues. Understanding the transcriptional networks and proteins that maintain appropriate pancreatic β cell maturation and function are essential for understanding the pathophysiology of diabetes to facilitate improved treatment options and/or leading to a cure for diabetes. In the pancreas, numerous studies over the past 30 years have identified many of the transcriptional regulatory pathways that regulate islet cell development and function (1–3). Each islet population is specified and maintained by distinct transcriptional programs that help establish its proper functional identity.

NKX2.2 is an essential transcription factor required for the proper development, maturation and function of β cells in mice and humans (4–7). NKX2.2 has 3 distinct protein domains: the tinman (TN) domain, the homeodomain (HD) which binds DNA and the NK2 specific (SD) domain. Mice lacking the TN domain (*Nkx2.2^TNMut^*) resulted in dysfunctional β cells that were either immature or transdifferentiated into α cells (8). Molecular experiments showed the NKX2.2 TN domain is required for the repressive activity of NKX2.2 by recruiting the Transducer-like enhancer of split 3 (TLE3)/Groucho-related gene 3 (GRG3) and Histone deacetylase 1 (HDAC1) to NKX2.2 repressed targets, including the master α cell regulator Aristaless homeobox gene *(Arx)* promoter (8). Characterization of mice lacking the SD domain (*Nkx2.2^SDMut^*) resulted in elevated blood glucose levels due to down regulation of β cell maturation and identity genes, among other phenotypes (9). Co-factors that interact with the SD domain have yet to be identified.

Although these previous studies identified functional roles associated with each of the NKX2.2 protein domains, many of the molecular mechanisms associated with each domain remain unknown. To address these gaps in knowledge, we set out to uncover essential cofactor proteins of NKX2.2 and determine their role in β cells. This analysis identified the Nucleosome Remodeling and Deacetylase (NuRD) complex and specifically the Chromodomain Helicase DNA-binding Protein 4 (CHD4) as NKX2.2 interacting factors. The generation of mice that deleted *Chd4* specifically in the developing β cells demonstrated that CHD4 is important for the structural integrity of islets and the proper maturation and function of β cells.

## Methods

### Identification of NKX2.2 interacting factors

Mouse Insulinoma (MIN6) β cell lines (10) were forward transfected with each of the *Nkx2.2* mutant plasmids or full-length *Chd4* and empty vector controls according to standard protocols. The myc epitope tag was used to pull down the respective NKX2.2 proteins and anti-CHD4 Ab was used to pull down endogenous CHD4. Lysates were collected for MS and western analyses according to standard protocols. Extensive experimental details are provided in supplemental methods.

### Generation of Chd4 KO Mice and Animal Maintenance

*Chd4* pancreas KO mutant mice were created by breeding *Pdx1^(Cre/+)^*mice (B6.FVB-Tg(Pdx1-cre)6Tuv/J) (11) and *Chd4^(flox/flox)^*(Chd4^tm1.1Kge^) created in the Georgopoulos lab (12)*. Chd4* βKO mutant mice were created by breeding *Ins1^(Cre/+)^* mice (B6(Cg)-*Ins1^tm1.1(cre)Thor^*/J) (13) and *Chd4^(flox/flox)^* mice. For experiments requiring cell sorting, *Ins2^(GFP/+)^* reporter mice (14) were bred into the *Chd4* control and βKO line. Physiological studies were conducted according to standard procedures on male and female mice with ages of mice being noted for each experiment.

Experimental details provided in supplemental methods. Mice were kept in accordance with University of Colorado Institutional Animal Care and Use Committee (IACUC) protocol #00045. Genotyping primers are listed in Supplementary Table 1.

### Immunofluorescence Staining and Image Analysis

Mouse pancreata were dissected and processed according to standard protocols. Every 10^th^ slide, spanning the entire pancreas was stained and analyzed. Full islet area was quantified as well as positive hormone area by quantifying all area above a set threshold. Hormone area for P2 was normalized by total pancreas area and hormone area for 3- and 6-week timepoints was normalized by total islet area.

### Glucose Stimulated Hormone Secretion Assays and Analysis

Pancreas tissue slices were prepared as previously described (15). Slices were used to perform GSIS and intracellular calcium imaging experiments. Insulin was assayed using static hormone secretion assays. For glucose stimulation measurements, each slice set was placed under a sequential 30-minute stimulation condition under one of three conditions: (1) 2 mM glucose, (2) 11 mM glucose, or (3) 20 mM glucose. For GIRK inhibitor measurements 20 mM glucose was supplemented with either 20 mM KCl or 10 µM VU0468554 (selective GIRK inhibitor; Axon Medchem, Cat. #3593). Insulin concentrations were measured using a mouse ultrasensitive insulin ELISA kit (Crystal Chem, Cat. #90096) per the manufacturer’s instructions. Calcium imaging and analysis is described in supplemental methods.

### Statistical Analysis

Unless otherwise stated, error bars are shown as mean +/- standard deviation (SD) and Shapiro-Wilk test was used to test normality. If passed, unpaired two-tailed student’s t-test was used to determine significant differences, otherwise, Mann-Whitney U test was used. P-value or padj of ≤ 0.05 was used to determine statistical significance. If no statistics are shown on graphs, results were not significant.

### Data and Resource Availability

#### Data availability

All data generated or analyzed during this study are included in the published article and its online supplementary files.

#### Resource availability

No novel resources were generated in this study; however, all resources are available from the corresponding author upon request.

## Results

### CHD4 and NuRD complex proteins interact with NKX2.2 in pancreatic β cells

NKX2.2 functions as a repressor and activator in pancreatic islets (4; 6; 8; 9). To identify cofactors of NKX2.2 that may influence its function in pancreatic β cells, we used co-IP followed by mass spectrophotometry (MS) using the full-length myc-tagged NKX2.2 protein and several previously characterized *Nkx2.2* mutant derivatives (TN^mut^, SD^mut^, TN^mut^/SD^mut^) (8; 9) (Fig. 1A). The immunoprecipitation with NKX2.2 wild type (WT) protein identified the largest number of interacting proteins, with 211 proteins enriched at least 2-fold in abundance compared to the empty vector (EV) control. The NKX2.2 TN^mut^ and NKX2.2 SD^mut^ protein derivatives identified 169 and 143 interacting proteins, respectively, that were enriched at least 2-fold in abundance compared to the EV control. Interestingly, the NKX2.2 TN^mut^/SD^mut^ mutant derivative only identified ∼84 interacting proteins, suggesting that deletion of the SD domain disrupted a substantial number of interactions, independently and in cooperation with the TN domain.

**Figure. 1.**
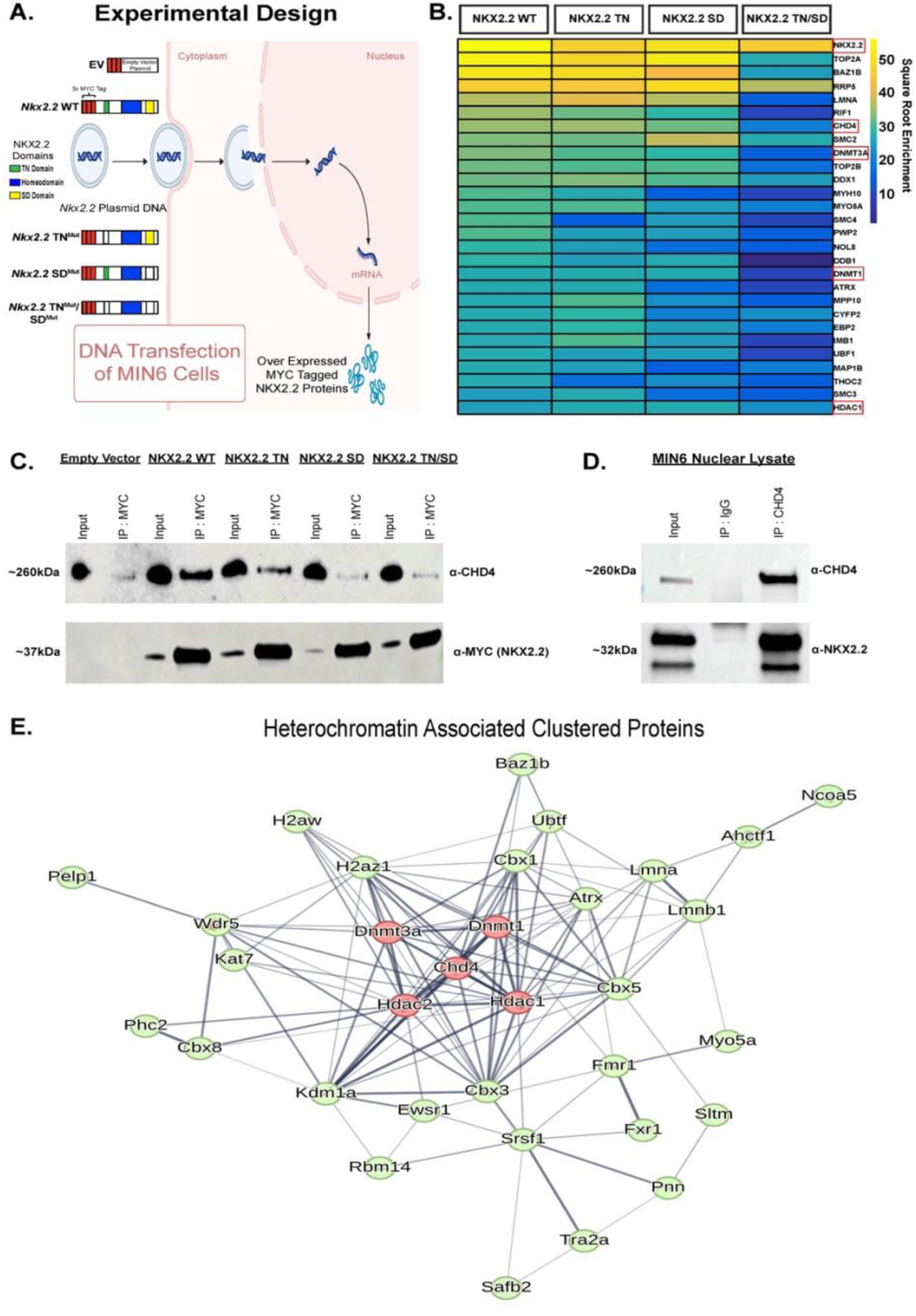
NKX2.2 interacts with chromatin modifying protein CHD4. (A) Schematic for transfection of *Nkx2.2* plasmids with WT and mutant domains into MIN6 β cell culture line for IP-MS. Empty Vector (EV) used as control. (B) Heatmap showing a subset of the highest enriched proteins to come out of the NKX2.2 IP-MS for WT NKX2.2 and each domain mutant. Color represents square root normalization of enrichment. (C) Western blot of MYC tag IP in MIN6 nuclear lysate. (D) Western blot of CHD4 IP in MIN6 nuclear lysate. (E) Heterochromatin group cluster from string database clustering (23) showing overlapping CHD4 and NKX2.2 interacting proteins.

In addition to identifying known NKX2.2 interacting proteins, including DNMT3A, DNMT1 and HDAC1 (8; 16), several members of the Nucleosome Remodeling and Deacetylase (NuRD) complex, including the functional enzymatic protein Chromodomain Helicase DNA-binding Protein 4 (CHD4), were identified in the screen (Fig. 1B). CHD4 showed the highest enrichment in the NKX2.2 WT and NKX2.2 TN^mut^ samples and the interaction appeared to be weakened in the NKX2.2 SD^mut^ and NKX2.2 TN^mut^/SD^mut^ samples. Validation of the interaction between NKX2.2 domains and CHD4 showed a similar pattern of enrichment observed in the IP-MS (Fig. 1C). Interestingly, the interaction between NKX2.2 and CHD4 was disrupted to a greater extent in the NKX2.2 SD^mut^ and NKX2.2 TN^mut^/SD^mut^ samples, suggesting the SD region may be the main domain facilitating the interaction of NKX2.2 and CHD4.

Both the NuRD complex and CHD4 specifically are necessary for the development, maturation and function of many different tissues and cells, often acting as gene repressors by deacetylating histones and remodeling chromatin (17–21), although activation of genes has been noted as well (12; 22). To further validate an interaction between CHD4 and NKX2.2, we immunoprecipitated CHD4 from MIN6 nuclear lysate followed by MS. As expected, this identified the other NuRD complex components (HDAC1 and 2, MTA1, 2 and 3, RBBP 4 and 7, MBD 2 and 3), many of which were also present in the NKX2.2 WT MS. NKX2.2 was also identified in the CHD4 IP-MS and was verified by pulling down CHD4 in MIN6 nuclear lysate and western blotting for endogenous NKX2.2 (Fig. 1D).

Comparison between proteins that interacted with either NKX2.2 WT and CHD4 identified 112 proteins as cofactors of both NKX2.2 and CHD4. Greater than half of the significantly enriched protein cofactors identified in the NKX2.2 WT MS were also significant CHD4 protein cofactors and a hypergeometric test confirmed that this represents a statistically significant enrichment (p < 0.001). A String database (23) was used to group the 112 proteins using K means clustering into 3 main clusters (Supplementary Fig. 1A). CHD4 and 32 other proteins were clustered as a protein group associated with heterochromatin. Interestingly, in addition to HDAC1 and HDAC2, DNMT3A and DNMT1 interacted with both CHD4 and NKX2.2 (Fig. 1E). Due to the known repressor functions of DNA methyltransferases and histone deacetylases (24), we hypothesized that NKX2.2 recruits CHD4 to specific sites in the β cell genome to facilitate repression by closing chromatin and repressing non-β cell genes.

### Pancreas knock out of Chd4 leads to blood glucose and body weight defects

*Chd4* is expressed in all pancreatic islet cells types (25) and has a role in regulating adult β cell function through an interaction with Pancreas Duodenum Homeobox1 (PDX1) (26; 27); however, its role in regulating islet development and maturation, where NKX2.2 plays a major role, has not yet been assessed. To determine whether CHD4 functions during pancreas development we generated mice carrying a *Chd4* floxed (*Chd4^(fl/fl)^)* (12) allele and a *Pdx1^(Cre/+)^* (11) allele which would remove CHD4 from the entire pancreas and duodenum at the onset of pancreas formation (Supplementary Fig. 1B). The *Chd4^(fl/fl)^; Pdx1^(Cre/+)^* mice displayed a significant increase in the *ab libitum* blood glucose as early as postnatal day 2 (P2) and through 4 weeks of age compared to *Chd4^(fl/fl)^* and *Chd4^(fl/+)^; Pdx1^(Cre/+)^* control mice (Supplementary Fig. 1C). Morphometric analysis did not reveal an alteration of islet cell numbers or islet architecture at P2; but there was a severe disruption in islet structure at 4 weeks of age (Supplementary Fig. 1D). However, there was also a significant decrease in body weight of mutant *Chd4^(fl/fl)^; Pdx1^(Cre/+)^* mice compared to *Chd4^(fl/fl)^* and *Chd4^(fl/+)^; Pdx1^(Cre/+)^* mice at P2 and 4 weeks of age (Supplementary Fig. 1C). This suggests that removing *Chd4* from the pancreatic exocrine, ductal, and intestinal duodenal cells, in addition to loss of *Chd4* from the pancreatic endocrine lineage may disrupt the proper secretion of digestive enzymes and/or digestion of foods. Since assessment of endocrine pancreas function could be confounded by the defects in these other tissues that caused significant body weight loss, we chose to focus on the role of CHD4 specifically in the developing β cells.

### Chd4 βKO does not cause an overt developmental phenotype

To characterize the function of CHD4 in developing β cells from the onset of their differentiation, we generated *Chd4^(fl/fl)^; Ins1^(Cre/+)^* (13) mice (hereafter referred to as *Chd4* βΚΟ mice). Deletion of the CHD4 protein specifically from the β cell lineage was confirmed through Western blot analysis and IF staining (Supplementary Fig. 2A and 2B). At all ages tested, deletion of *Chd4* from the β cells did not result in a body weight phenotype (Supplementary Fig. 2C). To characterize *Chd4* βKO mice, we performed morphometric analysis and assessed blood glucose at postnatal day 2 (P2). Unexpectedly, given the severe developmental phenotypes associated with deletion of *Pdx1* or *Nkx2.2* in developing β cells (4; 7; 28-30), and the known interaction of PDX1 and NKX2.2 with CHD4, the neonatal *Chd4* βKO mice had normal blood glucose levels (Fig. 2A), as well as no change in the hormone area of insulin, glucagon or somatostatin (Fig. 2B), compared to control mice. There were also no apparent alterations in the islet architecture or structure when comparing *Chd4* βKO mouse islets to control mouse islets (Fig. 2C), suggesting that CHD4 is not necessary for specification or early development of the β cell lineage.

**Figure. 2.**
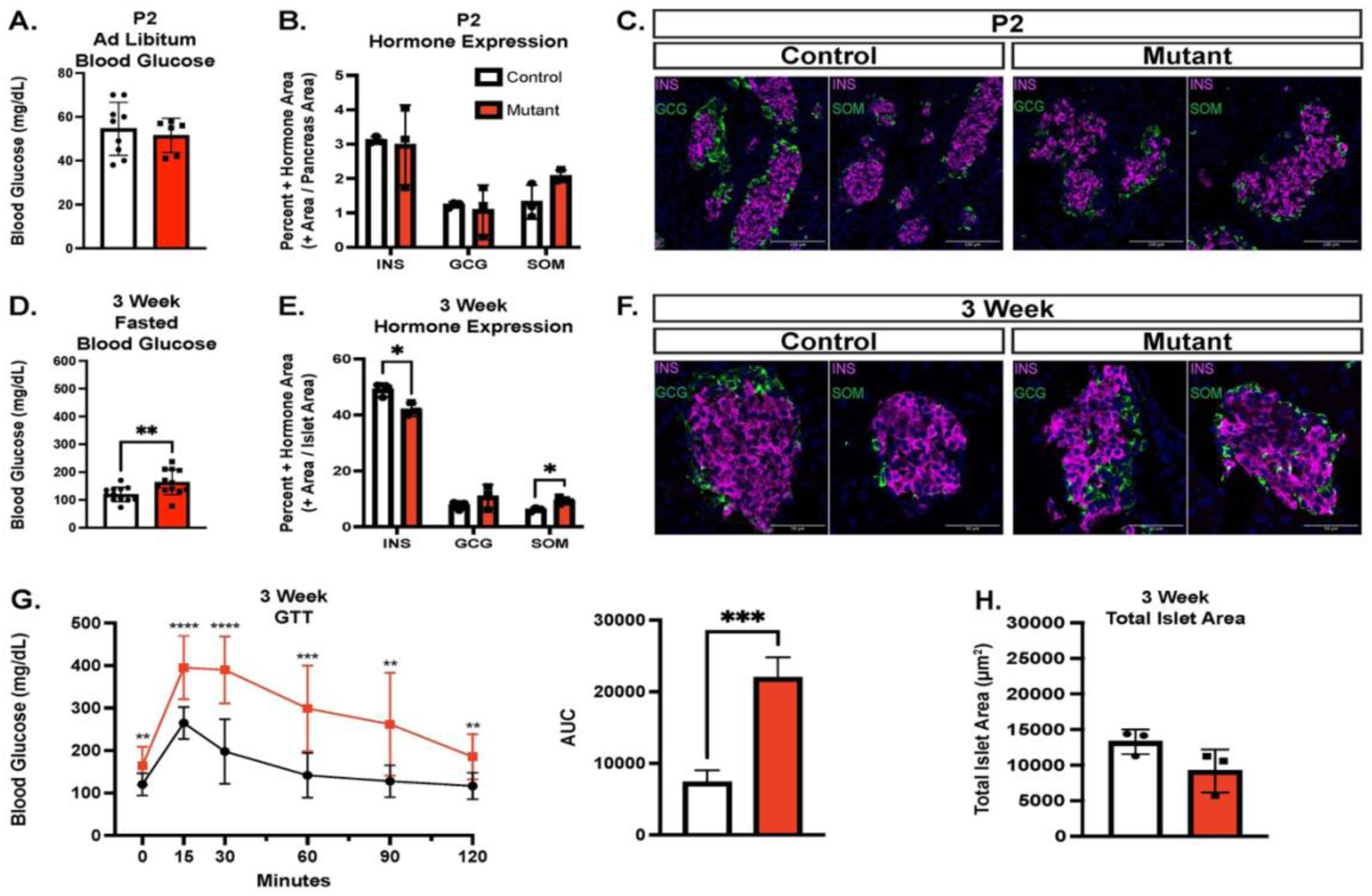
*Chd4* βKO mice develop phenotype at 3 weeks with hyperglycemia and disrupted islet architecture. Unless otherwise noted, mouse data consists of control (*Chd4^fl/fl^* : white bars) and mutant/*Chd4* βΚΟ (*Chd4^fl/fl^; Ins1^Cre/+^*: red bars) genotypes. (A) Measurements of *Ad libitum* blood glucose for P2 control (n =6) and mutant (n=9) mice. (B) Islet hormone quantification at P2 for Insulin (INS), Glucagon (GCG) and Somatostatin (SOM) for control (n=3) and mutant (n=3) mice. (C) Representative immunofluorescent images (20x magnification) for P2 control and mutant mice showing DAPI (blue), Insulin (INS, magenta), and Glucagon (GCG) or Somatostatin (SOM), green. Scale bars: 100 µm. (D) Measurements of fasted blood glucose for 3-week control (n =12) and mutant (n=12) mice. (E) Islet hormone quantification at 3-weeks for Insulin (INS), Glucagon (GCG) and Somatostatin (SOM) for Control (n=3) and Mutant (n=3) mice. (F) Representative immunofluorescent images (40x magnification) for 3-week control and mutant mice showing DAPI (blue), Insulin (INS, magenta), and Glucagon (GCG) or Somatostatin (SOM), green. Scale bars: 50 µm. (G) Glucose Tolerance Test (GTT) of 3-week control (black line, n=12) and mutant (red line, n=12) mice (multiple comparison two-tailed Student’s *t*-test). Area under the curve (AUC) measurement for GTT. All points normalized to 0 minutes time point prior to AUC calculation. (H) Quantification of 3-week control (n=3) and mutant (n=3) total islet area. For all figures, \**P*≤0.05, \*\**P*≤0.01, \*\*\**P*≤0.001, \*\*\*\**P*≤0.0001; two-tailed Student’s *t*-test unless otherwise specified.

### Chd4 βKO mice are glucose intolerant and hyperglycemic by 3 weeks of age

At 3 weeks of age, combined male and female *Chd4* βΚΟ mice had significantly higher fasting blood glucose levels (Fig. 2D) when compared to control mice that included *Chd4^(fl/fl)^, Ins1^(Cre/+)^*(*Cre* only control) and *Chd4^(fl/+)^; Ins1^(Cre/+)^* (Het) mice. Because none of the control mice displayed defects in fasting blood glucose levels or glucose tolerance (Supplementary Fig. 3), subsequent experiments included only *Chd4^(fl/fl)^*(control) and *Chd4^(fl/fl)^*; *Ins1^(Cre/+)^* (*Chd4* βKO) mice. When assessing sex separately at 3 weeks, only female *Chd4* βKO mice showed significantly higher fasting and *ad libitum* blood glucose levels compared to controls and were glucose intolerant (Supplementary Fig. 4A). Morphometric analysis performed on combined male and female mice at 3 weeks of age showed a significant decrease in the insulin positive hormone area, no change in the glucagon positive area, and a slight but significant increase of the somatostatin positive hormone area in *Chd4* βΚΟ compared to control mice (Fig. 2E). In addition to the changes in hormone expression area, islet architecture was disrupted in *Chd4* βΚΟ mice compared to control mice; glucagon and somatostatin expressing cells, typically present at the periphery of the islet, were present in the islet core where only β cells are normally seen (Fig. 2F). Interestingly, we did not observe the formation of polyhormonal cells, as was seen in the Nkx2.2 βKO mice (6). Lastly, combined male and female *Chd4* βΚΟ mice were significantly more glucose intolerant at 3 weeks (Fig. 2G) when compared to control mice.

Interestingly, the significant decrease in the insulin positive hormone area at 3 weeks was not accompanied by a change in the total islet area between *Chd4* βKO and control mice (Fig. 2H).

### Chd4 βKO mice have reduced insulin content at 6 weeks of age and are overtly diabetic by 10 weeks of age

By 6 weeks of age, fasting and *ad libitum* blood glucose levels in both the male and female *Chd4* βΚΟ mice were significantly elevated compared to controls (Fig. 3A, Supplementary Fig. 4B). There was still a significant decrease in the insulin positive hormone area between *Chd4* βΚΟ mice and control mice, while both glucagon and somatostatin positive hormone areas were unchanged (Fig. 3B). At this age, islet architecture continued to be disrupted with peripheral glucagon and somatostatin expressing cells present in the interior of the islets, but there was still no evidence of polyhormonal cells (Fig. 3C). Also at 6 weeks, the *Chd4* βΚΟ mice became progressively more glucose intolerant (Fig. 3D), though the significant decrease in the insulin positive hormone area at 6 weeks was still not accompanied by a change in the total islet area between *Chd4* βKO and control mice (Fig. 3E), suggesting β cells were maintained, but not expressing insulin.

**Figure. 3.**
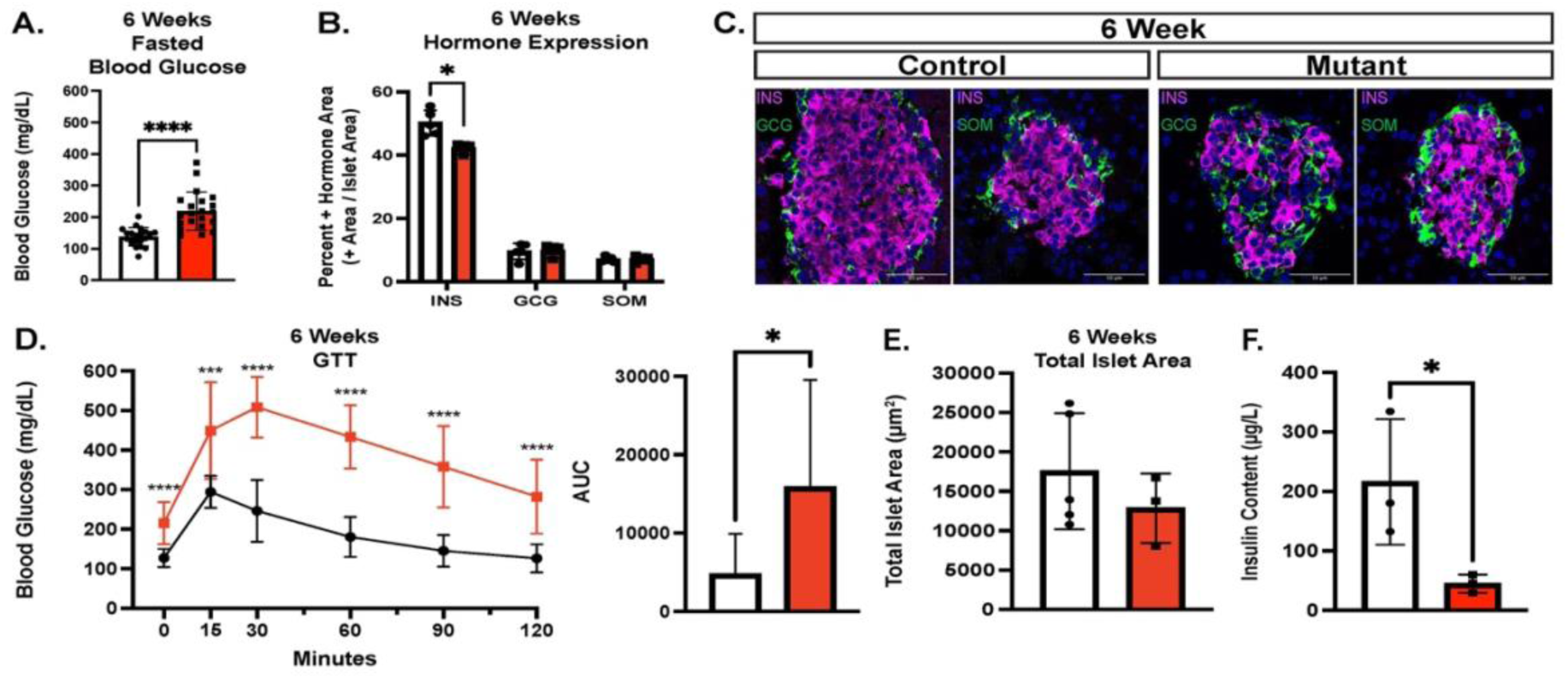
*Chd4* βKO mice remain hyperglycemic and have dysfunctional β cells. Unless otherwise noted, mouse data consists of control (*Chd4^fl/fl^* : white bars) and mutant/*Chd4* βΚΟ (*Chd4^fl/fl^; Ins1^Cre/+^*: red bars) genotypes. (A) Measurements of fasted blood glucose for 6-week control (n =20) and mutant (n=23) mice. (B) Islet hormone quantification at 6-weeks for Insulin (INS), Glucagon (GCG) and Somatostatin (SOM) for control (n=5) and mutant (n=3) mice. (C) Representative immunofluorescent images (40x magnification) for 6-week control and mutant mice showing DAPI (blue), Insulin (INS, magenta), and Glucagon (GCG) or Somatostatin (SOM), green. Scale bars: 50 µm. (D) Glucose Tolerance Test (GTT) of 6-week control (black line, n=13) and mutant (red line, n=13) mice (multiple two-tailed Student’s *t*-test). Area under the curve (AUC) measurement for GTT. All points normalized to 0 minutes time point prior to AUC calculation. (E) Quantification of 6-week control (n=5) and mutant (n=3) total islet area. (F) Insulin content from dissected mouse pancreas of 6-week Control (n=3) and Mutant (n=3). For all figures, \**P*≤0.05, \*\*\**P*≤0.001, \*\*\*\**P*≤0.0001; two-tailed Student’s *t*-test unless otherwise specified.

By 10 weeks of age, the *Chd4* βΚΟ male mice became severely diabetic compared to controls, displaying significantly lower body weight, significantly higher fasted and *ad libitum* blood glucose levels, severe glucose intolerance (Supplementary Fig. 4C) and dehydration due to excessive urination. The hyperglycemic phenotype and health of the female mice also continued to worsen through 10 weeks of age (Supplementary Fig. 4C), although the decline progressed more slowly. Further phenotypic and molecular analysis focused on the 3 week and 6 week timepoints to minimize secondary affects associated with the overt diabetic phenotype.

To analyze the functionality of the *Chd4 β*KO β cells we attempted to isolate islets to perform GSIS and calcium imaging, however, the *Chd4 β*KO mutant islets rapidly dissociated upon manipulation, revealing a striking islet fragility phenotype. To initially circumvent this challenge, insulin content from intact whole pancreata isolated from 6-week-old mice was measured. *Chd4* βΚΟ mice displayed significantly less insulin content compared to control mice (Fig. 3F). We therefore wanted to explore the transcriptional regulation in *Chd4 β*KO mice that may be contributing to the observed diabetic phenotypes.

### Chd4 mutant β cells have an immature transcriptional landscape

With CHD4’s role as a chromatin remodeling factor, it was important to identify the transcriptional defects that could explain the observed β cell dysfunction. To purify the β cell population we introduced an *Ins2^(GFP/+)^*allele (14) into the control and *Chd4* βΚΟ mice for Fluorescence Activated Cell Sorting (FACS) (Supplementary Fig. 5). Transcriptome analysis on β cells isolated from 4-week-old mice revealed many genes significantly dysregulated in the *Chd4* βΚΟ mice vs. controls (Fig. 4A), including a significant decrease in many essential β cell genes/factors including *Ucn3, MafA* and *Foxo1.* There was also a reduction in many of the genes involved in glucose sensing and insulin secretion including *Slc2a2* (GLUT2)*, Syt13, Ero1lβ, G6pc2*, *Slc30a8* (ZnT8), and *Glp1r* (Fig. 4B). Interestingly, *Chd4* βΚΟ mice also displayed a significant increase in genes normally expressed in immature β cells including *MafB* and *Npy* or pancreas disallowed genes such as *Hk1*. We also discovered *Kcnj5,* which encodes the GIRK4 channel that is normally not expressed in pancreatic β cells, as upregulated. GIRK4 is known to respond to somatostatin and epinephrine signaling (31; 32). Consistent with the lack of polyhormonal cells in the IF analysis, there was not a corresponding increase in non-β endocrine cell genes. Lastly, we also observed significant downregulation of *Robo2* (Fig. 4B). Previous studies demonstrated that deletion of *Robo1* and *Robo2* from β cells (*Robo* βKO) caused islets to easily dissociate (33; 34). Furthermore, deletion of *Robo2* alone from the β cells was sufficient to disrupt the islet architecture (35; 36), similar to the phenotypes observed in the *Chd4* βΚΟ mice (Figs. 2F, 3C). Overall, transcriptional analysis of the *Chd4* βKO β cells identified disruption of many β cell maturity and functional markers suggesting this was the underlying cause of the severe CHD4 mutant phenotype.

**Figure. 4.**
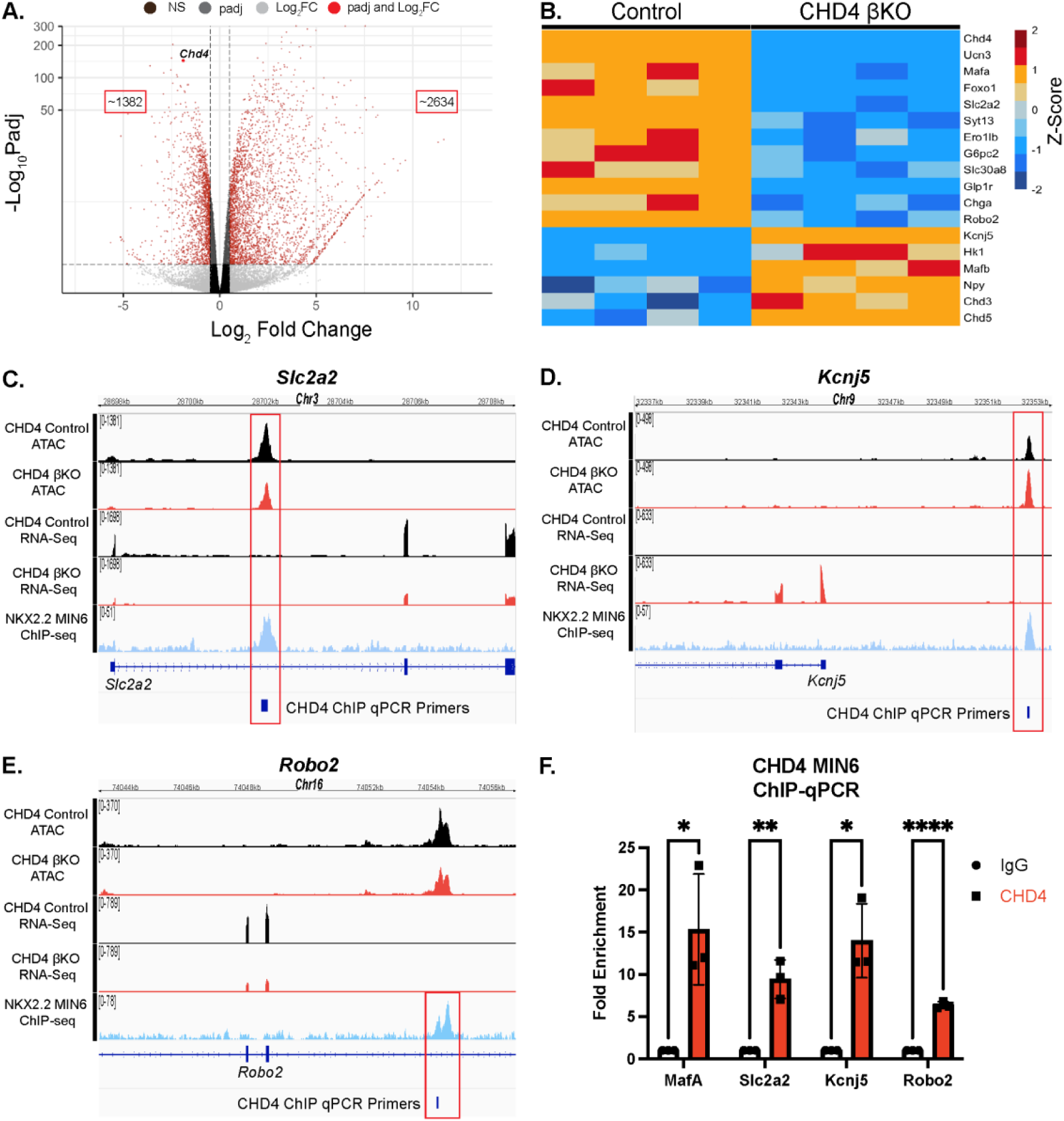
CHD4 binds and regulates essential pancreatic β cell genes. (A) Volcano plot showing the distribution of differentially expressed genes (DEGs) in *Chd4* βKO mice. (B) Heatmap showing a subset of differentially expressed β cell genes between control and *Chd4* βKO mice. Heat map is mean centered using z-score scale. (C-E) IGV tracks showing genome locations and sequencing read peaks of *Chd4* control and βKO ATAC-seq, *Chd4* control and βKO RNA-seq and *Nkx2.2* ChIP-seq datasets overlapped with CHD4 binding sites through qPCR primer locations for *Slc2a2, Kcnj5* and *Robo2.* (E) Quantification of qPCR using fold enrichment over IgG of *MafA, Slc2a2, Kcnj5* and *Robo2* representing binding sites of CHD4 at these genes (two-tailed Student’s *t*-test). \**P*≤0.05, \*\**P*≤0.01, \*\*\*\**P*≤0.0001.

### CHD4 affects NKX2.2 - associated β cell gene regulation through chromatin remodeling

The identification of CHD4 as an NKX2.2 co-factor suggested there would be a set of genes that were co-regulated by NKX2.2 and CHD4 and were associated with changes in the chromatin landscape when CHD4 was deleted. To detect changes in chromatin landscape, we reanalyzed a previously published CHD4 Control ATAC-seq vs. adult CHD4 βKO ATAC-seq dataset (27).

As expected for a chromatin modifier protein, there were greater than 52,500 total peaks of significantly altered chromatin (padj < 0.05) when CHD4 was disrupted. The majority of peaks (39,956) displayed regions of increased open chromatin, although there were many regions (12,843) that became increasingly closed. To identify genes that are potentially direct targets of NKX2.2 and CHD4, we assessed genes that were bound by NKX2.2, dysregulated in both the *Nkx2.2* βKO and *Chd4* βKO islets and displayed changes in chromatin accessibility. 62 genes were present in all five datasets (Supplementary Fig. 6A) of which 40 were dysregulated in the same direction in both the *Nkx2.2* βKO and *Chd4* βKO RNAseq datasets (Supplementary Fig. 6B), including *Kcnj5, Ucn3, Slc2a2, Slc30a8, G6pc2 and Npy.* These genes represent potential direct targets of CHD4 and NKX2.2 and have the potential to contribute to the observed *Chd4* βKO phenotype.

### β cell maturation genes are direct targets of NKX2.2 and CHD4

A subset of the potential NKX2.2 direct gene targets that were also regulated by CHD4 correlated with the observed *Chd4* βKO diabetic and fragile islet phenotypes. *Slc2a2,* the gene encoding the β cell glucose transporter GLUT2 is bound by NKX2.2, displayed reduced chromatin accessibility in the CHD4 βKO ATAC samples and was significantly down regulated when either *Nkx2.2* or *Chd4* were deleted from β cells (Fig. 4C). Alternatively, *Kcnj5,* which encodes the GIRK4 channel, is bound by NKX2.2, had increased chromatin accessibility in the CHD4 βKO samples, and was up regulated when either *Nkx2.2* or *Chd4* were deleted from β cells (Fig. 4D). Dysregulation of either gene could directly contribute to the observed insulin secretion defects. In addition, we assessed whether CHD4 directly regulated *Robo2*, to potentially contribute to the observed fragile islet phenotype (Fig. 4E, 2F, 3C).

Chromatin immunoprecipitation of CHD4 bound regions followed by qPCR (ChIP-qPCR) in MIN6 cells demonstrated that CHD4 binding was significantly enriched at *Slc2a2, Kcnj5* and *Robo2* loci compared to IgG (Fig. 4F), suggesting each of these genes as direct targets of CHD4. The *MafA* regulatory region 3 region, which contains a characterized CHD4 binding site (26), was included as a positive control for these experiments. Interestingly, this analysis of CHD4 direct targets suggested that while CHD4 is necessary to close chromatin for many genes, such as *Kcnj5*, it also appeared to be opening chromatin at several important β cell genes, such as *Slc2a2*.

### Chd4 mutant β cells show disrupted calcium signaling

We next tested whether the function of the *Chd4* βΚΟ islets was impaired, as suggested by the diabetic phenotype and by the differential gene expression and chromatin accessibility observed in the mutant animals. With the observed up regulation of *Kcnj5* in the *Chd4* βΚΟ islets, and previous research showing that GIRK channels may disrupt islet cell polarization (31), we first tested whether the *Chd4* βKO mice had disrupted calcium signaling. To circumvent the islet fragility phenotype, we employed live pancreas slices to assess the status of calcium signaling in *Chd4* βΚΟ islets (15). Using the Ca^2+^ sensor, Fluo-4, we measured the Ca^2+^ response in control and *Chd4* βΚΟ pancreatic slices upon elevated glucose. This analysis revealed that islets within *Chd4* βΚΟ slices displayed a severely impaired Ca^2+^ response to elevated glucose, including the absence of an initial elevation in Ca^2+^, and lack of pulsatile waves of Ca^2+^ compared to control islet slices (Fig. 5A and B, Supplementary Movie 1 and 2). The mean Ca^2+^ elevation, expressed as the mean fluorescence increase from 0-5 minutes and 5-25 minutes each had significantly lower signal in the *Chd4* βΚΟ slices than in the control slices (Figs. 5C, 5D). These experiments suggest our *Chd4* βKO islets have disrupted β cell calcium signaling, which could severely impair the β cell insulin secretion.

**Figure. 5.**
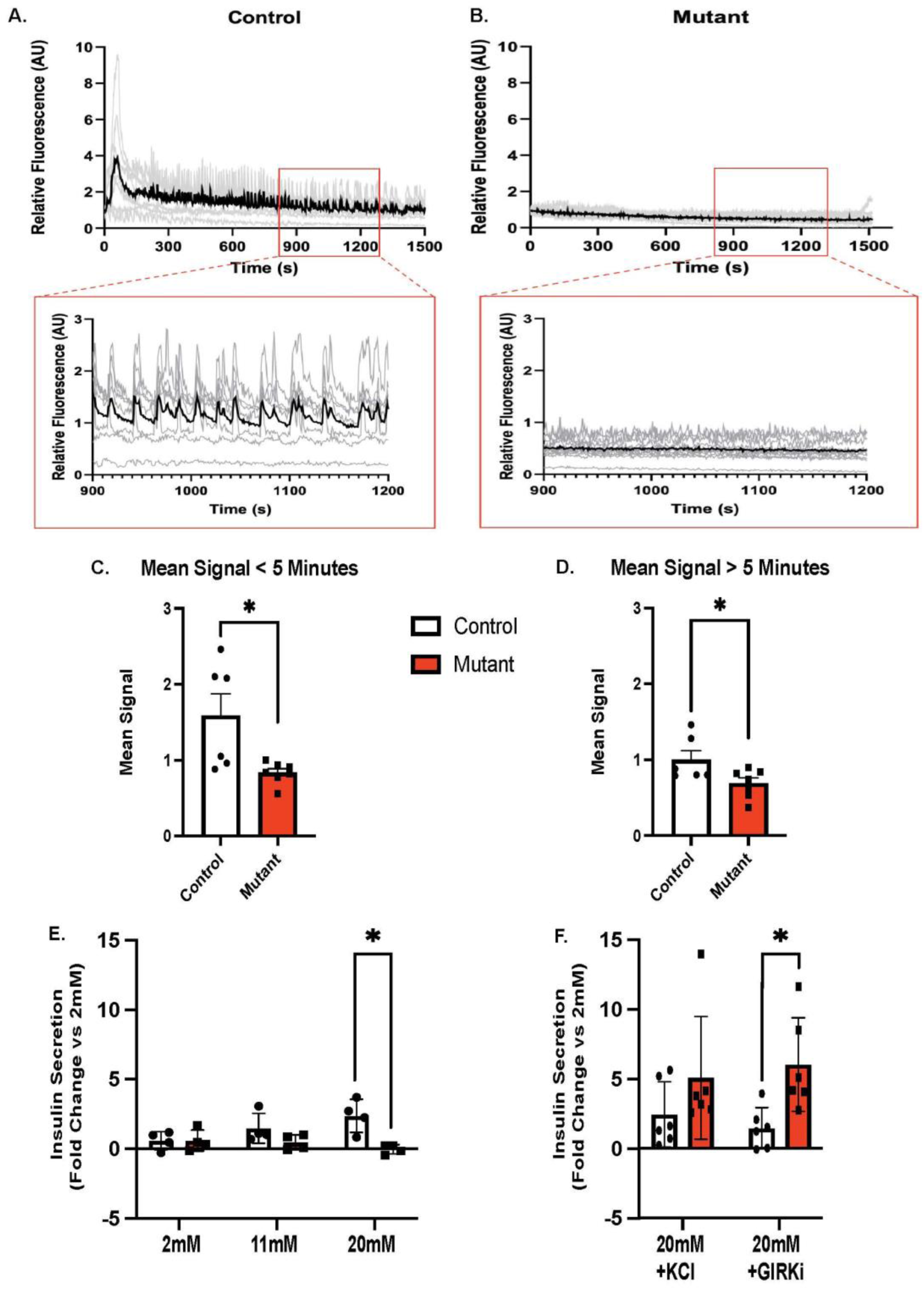
Calcium signaling is disrupted in *Chd4* βKO pancreas slice culture islets. (A) Time-course of the Fluo4 Ca^2+^ sensor fluorescence for one islet from a control pancreas slice following elevation of glucose to 11mM. Each gray trace is one cell from the islet, and the black trace is the average fluorescence of all cells from the islet. Fluorescence is normalized to the mean fluorescence at 2mM glucose. Inset box is zoomed in view from 900 to 1200 seconds showing coordinated calcium oscillations in the control islet. (B) Time-course of the Fluo4 Ca^2+^ sensor fluorescence for one islet from mutant pancreas slice, as in A. (C) Quantification of mean Fluo4 signal for 0 minutes to 5 minutes after Ca^2+^ elevation, for control (n=6) and mutant (n=6) pancreas slices. Mean signal = average fluorescence over indicated time (two-tailed Student’s *t*-test). (D) Quantification of mean signal for 5 minutes to 25 minutes after Ca^2+^ elevation for control (n=6) and mutant (n=6) pancreas slices, as in C (two-tailed Student’s *t*-test). (E) Glucose Stimulated Insulin Secretion (GSIS) assay of 6-week control and mutant pancreas slices. Conditions were 2mM glucose, control (n=4) and mutant (n=4), 11mM glucose, Control (n=4) and Mutant (n=4), 20mM glucose, control (n=4) and mutant (n=4). (F) Glucose Stimulated Insulin Secretion (GSIS) assay of 6-week control and mutant pancreas slices. Conditions were 20mM glucose + KCl, control (n=6) and mutant (n=6) and 20mM glucose + GIRK inhibitor (GIRKi, 10µM VU0468554), control (n=6) and mutant (n=6). GSIS is expressed as a fold-change for each condition normalized to the secretion measured in the same slices at 2mM glucose. For all figures, \**P*≤0.05; two-tailed Student’s *t*-test unless otherwise specified.

### Chd4 βKO mice have defective insulin secretion that can be rescued by inhibiting GIRK4

Consistent with the observed disruption of calcium signaling in the *Chd4* βΚΟ slices, Glucose Stimulated Insulin Secretion (GSIS) assays on the control vs. *Chd4* βΚΟ pancreatic slices demonstrated that insulin secretion from *Chd4* βΚΟ islets trended lower at 11mM glucose and was significantly lower at 20mM glucose (Fig. 5E), as compared to control islets. Treatment with 20mM glucose + KCl conditions restored insulin secretion from *Chd4* βKO slices (Fig. 5F), suggesting that acute depolarization of the β cell could bypass the calcium signaling defect. To investigate whether the upregulation of *Kcnj5* (GIRK4), was directly affecting insulin secretion, we used a GIRK channel inhibitor (VU0468554, GIRKi) to selectively inhibit GIRK1/GIRK4 channels (37) along with a 20mM glucose stimulation. The addition of the GIRKi at 20mM glucose showed a significant increase in insulin secretion in the *Chd4* βKO slices compared to controls (Fig. 5F), suggesting that the *Chd4* βΚΟ slices have a defect in glucose stimulated insulin secretion that can be rescued when GIRK4 is inhibited.

## Discussion

Prior studies in our lab have shown the essential role of NKX2.2 in the proper development, maturation and function of pancreatic β cells. Given its diverse functions, NKX2.2 likely interacts with co-factors to exert its activities. In this study, we identified about 200 proteins that potentially interact with NKX2.2 to facilitate transcriptional regulation in the β cell. Many of the interactions were mediated by either the TN domain, SD domain or both. This study identified novel interactions between NKX2.2 and several members of the Nucleosome Remodeling and Deacetylase (NuRD) complex, including CHD4, an enzymatic component of the complex, which appeared to be predominantly mediated by the NKX2.2 SD domain, although further studies would be needed to assess domain binding.

Although *Chd4* was deleted from β cells at the onset of insulin expression during embryogenesis, the *Chd4* βKO mice did not display an overt phenotype at P2. This result was somewhat surprising given its interaction with NKX2.2, which is essential for proper β cell development (4; 7). It is possible that *Chd4* is dispensable for embryonic β cell development. Alternatively, although we confirmed that CHD4 was efficiently removed from the majority of the adult β cells (Supplementary Fig. 2A, 2B), a small proportion of β cells that retained CHD4 expression could be sufficient to sustain glucose regulation through the early postnatal stages. Lastly, CHD4 function may also be compensated by other family members including CHD3 and CHD5, which were both upregulated in the *Chd4* βKO β cells (Fig 4B). However, even if CHD3 and CHD5 were able to compensate for CHD4 during pancreas development, given the striking severity of the *Chd4* βKO phenotype, these family members appear to be unable to compensate during postnatal β cell maturation and function.

Deletion of *Chd4* from the developing β cell lineage had a substantial impact on the maturation and function of β cells. *Chd4* βKO mice were glucose intolerant and hyperglycemic by 3 weeks of age with their symptoms worsening with age. Interestingly, a recent study that used a *MIP:Cre^ERT^*to remove *Chd4* from fully mature adult β cells (27) caused impaired insulin secretion; however, the phenotype was less severe than when *Chd4* was deleted from developing β cells, suggesting that in β cells CHD4 may play a stronger role in initiating chromatin modifications to establish a transcriptional program than maintaining chromatin structure. In addition, several important β cell transcription factors, including *Foxo1*, and β cell maturation factors such as *Ucn3* were not affected when *Chd4* was deleted from adult β cells, suggesting that maintenance of β cell identity and maturation is not dependent on CHD4. However, *Robo2* and *Slc2a2* gene transcription was disrupted in the *Chd4^fl/fl^; MIP:Cre^ERT^* mice, suggesting CHD4 is required to maintain β cell integrity and functional status.

This study begins to parse out novel roles of CHD4 in establishing the correct chromatin landscape and transcriptional status during the formation and maturation of β cells vs. its role in maintaining β cell functions. Davidson et al., (26) identified an interaction between CHD4 and PDX1 and showed the interaction of CHD4 facilitated by PDX1 may also be tuned through glucose sensing in the β cell. Although we failed to identify PDX1 as a CHD4 interacting factor in the MS analysis, the NKX2.2 MS did identify PDX1 as a potential NKX2.2 interacting factor. Consistently, NKX2.2 and PDX1 co-regulate many essential regulatory genes in the β cell genome including at *Mafa, Slc2a2 and Robo2,* although not all targets are shared, such as at *Kcnj5,* where only NKX2.2 binds (6; 38). This leads us to believe that a subset of CHD4 functions in the β cells are facilitated through its recruitment by both NKX2.2 and PDX1, while others may rely on either NKX2.2 or PDX1.

*Kcnj5* was one of the most upregulated genes in both *Chd4* βKO and *Nkx2.2* βKO RNAseq datasets (Supplementary Fig. 6B). *Kcnj5* encodes the G protein-activated inward rectifier potassium channel 4 (GIRK4). There is some discrepancy in the literature as to whether *Kcnj5* is normally expressed in mouse pancreatic β cells (25; 31; 32); however, it has been shown that somatostatin and epinephrine signaling can affect the GIRK4 channel, leading to potassium efflux from the cell, and hyperpolarizing the cell membrane of β cells and α cells (31; 32; 39-41). Consistent with this action, we observed Ca^2+^ activity and insulin secretion to be substantially disrupted by elevated glucose. Furthermore, glucose-stimulated insulin secretion upon either a GIRK1/4 channel inhibitor or KCl rescued the diminished insulin secretion at elevated glucose. This suggests that the GIRK4 channel upregulation in the *Chd4* βKO mice is at least in part driving the loss of insulin secretion and glucose control through aberrant hyperpolarization of the β cell membrane. However, we did not determine whether the downregulation of *Kcnj5* is the main driver of the disruption to the Ca^2+^ activity or acting in concert with other gene disruptions in the β cell. For example, the downregulation of *Slc2a2* (GLUT2) could reduce the amount of glucose entering the β cells, which would also cause a disruption in Ca^2+^ activity; this was not tested.

The identification of *Robo2* as significantly down regulated in the absence of Chd4, suggests that loss of *Robo2* may contribute to the fragile islet phenotype as well as the disrupted islet architecture and impaired calcium signaling (33; 35; 36). However, deletion of *Robo2* alone from β cells did not cause disruption of maturation and β cell identity markers (35), as observed in our study. This suggests the loss of β cell maturation and identity may be driven by direct targets of CHD4 such as *MafA* and *Slc2a2.* Furthermore, in addition to *Robo2*, multiple genes involved in cell-to-cell and cell-to-matrix junctions were also significantly down-regulated including many genes involved in desmosomes (*Jup, Pkp2, Dsc2*) which have been shown to affect tissue architecture and structure (42–44), suggesting additional CHD4 targets may contribute to the fragile islet and disrupted architecture phenotypes.

In other tissue and cell types, CHD4 has been shown to either activate genes, including those associated with lineage programs (12; 22), or repress genes associated with alternative lineage programs (20; 45). We demonstrated that in pancreatic β cells, CHD4 binding at specific loci correlated with both increased and decreased chromatin accessibility. We also observed a number of essential β cell genes were down regulated in the absence of *Chd4*; whereas genes such as *Kcjn5* were upregulated. However, since deletion of *Chd4* in β cells predominantly caused gene upregulation and increased chromatin accessibility, it is likely that CHD4 primarily functions as a repressor. In future studies, it would be interesting to identify whether the distinct NKX2.2 domains confer either NKX2.2 and/or CHD4 combined roles as repressors or activators in β cells.

Overall, understanding the transcriptional network that facilitates the proper development, maturation and function of pancreatic β cells can help lead to better treatment options for diseases like diabetes. This study increases our understanding of how essential transcription factors in the β cell, like NKX2.2, maintain a proper transcriptional state through chromatin modifications by interacting with cofactor proteins such as CHD4 to allow the proper maturation and function of β cells.

## Acknowledgments

We thank the Sussel lab for providing feedback about experimental design and concepts. A special thank you to Drs. David Lorberbaum and Fiona Docherty for critically reading the manuscript and providing feedback. We thank Dr. James Hagman for sharing the CHD4 mice and providing background information and advice about the NuRD complex. We also acknowledge Scott K. Beard from the Islet Isolation Core Facility, Barbara Davis Center for his specialty in isolating islets. This project was supported by funding from the National Institutes of Health, National Institute of Diabetes and Digestive and Kidney Diseases (NIDDK): Grants R01DK082590 (LS); R01DK106412 (RKPB); R01DK102950 (RKPB); R01DK140904 (RKPB); T32GM136444 (DKS); F31DK131639 (DKS) and the University of Colorado Diabetes Research Center (DRC) P30DK116073. Drs. Lori Sussel and Kristen Wells are the guarantors of this work and, as such, had full access to all the data in the study and take responsibility for the integrity of the data and the accuracy of the data analysis.

## Author contributions

Conceptualization: DKS, MAG, LS

Methodology: DKS, MAG, LS

Investigation: DKS, TGO, MAG, MRC, VMH, CRM

Data Analysis: DKS, CJH, KLW

Supervision: KLW, RKPB, LS

Writing—original draft: DKS, LS

Writing—review & editing: DKS, MAG, VMH, CJH, KLW, RKPB, LS

