## Supplemental Methods and Figures for "CHD4 and NKX2.2 Cooperate to Regulate Beta Cell Function by Repressing Non-Beta Cell Gene Programs"

##### *Cell Line and Cell Culture*

Mouse Insulinoma (MIN6) β cell culture line was originally created by the Yamamura lab (10) and cultured at 37°C and 5% $CO_2$. MIN6 cells were passaged every 7 days and fed media containing Dulbecco's Modified Eagle Medium (DMEM) high glucose, with L-glutamine, without Sodium pyruvate and HEPES (Gibco, Cat. # 11965-092) and supplemented with 10% FBS (Gibco, Cat. #A5669701), 5% PenStrep (10,000U/mL Penicillin-Streptomycin, Gibco, Cat. #15140-122) and 57 µM βME.

##### *Nkx2.2 Domain Variant Plasmid Transfections*

MIN6 cells were forward transfected with 31.5 µg of one of the following *Nkx2.2* plasmids: pcDNA3-3x Myc-EV, pcDNA3-3x Myc-Nkx2.2 WT, pcDNA3-3x Myc-Nkx2.2 $TN^{mut}$, pcDNA3-3x Myc-Nkx2.2 $SD^{mut}$, or pcDNA3-3x Myc-Nkx2.2 $SD^{mut}$ / $TN^{mut}$. Lipofectamine-3000 was used as the transfection reagent (Invitrogen, Cat. #L3000015) and MIN6 cells were transfected for 72 hours prior to protein extraction using the Nuclear Extract Kit (Active Motif, Cat. #40010). Protein concentration was assessed using the DC assay kit (Bio-Rad, Cat. #5000112).

##### *Co-Immunoprecipitation for Mass Spectrometry*

For mass spectrometry (mass spec) of transfected Myc tagged NKX2.2 proteins, pull downs were performed by incubating Anti-c-MYC magnetic beads (Pierce, Cat. #88843) with samples for 1 hour with mixing at 4°C. Samples were then washed with a Co-IP Buffer (50mM HEPES, 1mM EDTA, 5% glycerol, 0.5% Triton-X, 500nM NaCl). For mass spec of endogenous CHD4 protein, pulldowns were performed in WT MIN6 cells using the Nuclear Complex Co-IP Kit (Active Motif, Cat. #54001) per the manufacturer's instructions. Briefly, 500 µg of WT MIN6 nuclear extract was incubated in IP High Buffer (no detergent and no NaCl) using 5 µg of CHD4 rabbit antibody (D8B12, CST, Cat. #11912) or 5 µg of a control IgG rabbit antibody (CST, Cat. #2729), overnight with mixing. The second day, samples were incubated with Anti-Protein-A magnetic beads (Pierce, Cat. #88846) for 1 hour with mixing at 4°C. Samples were then washed using the IP High Buffer (no detergent and no NaCl) with and without BSA.

For both the NKX2.2 and CHD4 samples, 4x Laemmli buffer (Bio-rad, Cat. #1610747) plus βME was used as a sample buffer and samples were boiled for 5 minutes at 95°C and loaded in a Bio-Rad Mini-PROTEAN TGX Gel (4-20%, 10-well, 50 µL, Bio-rad, Cat. # 456-1094). Once all samples had collected into compact bands in the stacking gel portion, the gel was stopped and extracted and washed 3x for 5 minutes in Milli-Q water. The gel was stained in SimplyBlue SafeStain (Invitrogen, Cat. # LC6060) for 1 hour at RT. The gel was destained 2x for 1 hour in Milli-Q water. The gel bands were cut out and submitted to the mass spec core in microcentrifuge tubes filled with water for processing. pcDNA3-3x Myc-EV was used as the control sample in the NKX2.2 experiment and IgG was used as the control sample in the CHD4 experiment.

***NKX2.2 & CHD4 Mass Spectrometry***

Samples were tryptically digested prior to LC/MS/MS analysis according to an established protocol (11; 12). Briefly, under keratin-free conditions, stacking gel bands were excised from the gel and dehydrated before being incubated with 25 ng/µL of trypsin at 37°C for 18 hrs, after which the enzyme was deactivated with 10% formic acid (FA). Peptides were extracted from the gel fragments, dried via speedvac, and resuspended for mass spectrometric analysis. NKX2.2 samples were re-suspended in 20 µL of buffer and CHD4 samples were re-suspended in 14 µL of buffer. NKX2.2 samples were analyzed similar to a previous publication (13), with the following exceptions: MS/MS data was collected in positive ion polarity over mass ranges 260–1700 m/z at a scan rate of 8 spectra/second for MS scans and mass ranges 50–1700 m/z at a scan rate of 3 spectra/second for MS/MS scans.

2 µL of CHD4 samples were acquired on an Orbitrap Eclipse (Thermo Scientific) operated using intensity-dependent CID MS/MS to generate peptide ID’s and equipped with a Ultimate 3000 RSCLnano LC system (Thermo Scientific) using a previously published method with the following exceptions: The orbitrap MS1 scan mass range was set to 375-1500 m/z range and dynamic exclusion was set to 20 seconds instead of 14 seconds (14). PEAKS studio proteomics version 10.5 was used to extract, search, and summarize peptide identity results from IgG and CHD4 samples (PEAKS, Waterloo, ON, Canada). Peptide identifications were performed using the PEAKS DB search engine combined with PEAKS de novo sequencing. Extracted spectra were searched against the SwissProt Mus musculus database. The crude PEAKS studio proteomics extractor protein

abundance levels were used to compare relative abundances of proteins in each sample to determine protein interactors for CHD4.

***Co-Immunoprecipitation and Western Blotting***

For Co-IP and western of endogenous CHD4, pulldowns were performed in WT MIN6 cells using the Nuclear Complex Co-IP Kit (Active Motif, Cat. #54001) per the manufacturer's instructions. Briefly, 500 µg of WT MIN6 nuclear extract was incubated in IP High Buffer (no detergent and no NaCl) using 5 µg of CHD4 rabbit antibody (D4B7, CST, Cat. #12011) or 5 µg of a control IgG rabbit antibody (CST, Cat. #2729), overnight with mixing. The second day, samples were incubated with Anti-Protein-A magnetic beads (Pierce, Cat. #88846) for 1 hour with mixing at 4°C. Samples were then washed using the IP High Buffer (no detergent and no NaCl) with and without BSA. For all samples, including 5% input samples, 2x Laemmli buffer (Bio-rad, Cat. #1610737) plus βME was used as a sample buffer and samples were boiled for 5 minutes at 95°C and loaded in a Bio-Rad Mini-PROTEAN TGX Gel (4-20%, 10-well, 50 µL, Bio-rad, Cat. # 456-1094) with a Precision Plus Protein Ladder (Bio-rad, Cat. #1610375). Protein gels were transferred onto PVDF membranes and washed 3x for 5 minutes in TBS-T at RT. Membranes were blocked in TBS-T + 5% milk for 1 hour at room temp and membranes were washed 3x for 5 minutes in TBS-T at RT. Primary antibodies (Supplementary Table 2) were added at concentrations noted in TBS-T + 5% BSA overnight on a rocker at 4°C. The next day, membranes were washed 3x for 5 minutes in TBS-T at RT and an IP Veriblot HRP secondary antibody was added at 1:500 in TBS-T + 5% Milk (abcam, Cat. #ab131366). Membranes were washed 3x for 5 minutes in TBS-T at RT and imaged using SuperSignal West Pico PLUS Chemiluminescent Substrate (Thermo Scientific, Cat. #34580).

***Mouse Islet Western Blotting***

For western blot of endogenous CHD4 in mouse islets, islets from 3 mice of control or mutant genotype were isolated as described above and were pooled to form 1 control or mutant sample. Islets were lysed using 1x RIPA Buffer (Abcam, Cat. # 156034) + PIC tablets (Invitrogen, Cat. # A32961) for 30 minutes on ice. Samples were spun down at 12,000 rpm for 20 minutes at 4°C and protein concentration was assessed using the DC assay kit (Bio-Rad, Cat. #5000112). 50 µg of Control or Mutant Mouse Islet Lysate was used and western was run as described above. Vinculin was used as the loading control (Supplementary Table 2).

##### *Generation of Chd4 KO Mice and Animal Maintenance*

*Chd4* pancreas KO mutant mice were created by breeding *Pdx1$^{(Cre/+)}$* mice (B6.FVB-Tg(Pdx1-cre)6Tuv/J) (15) and *Chd4$^{(flox/flox)}$* (Chd4$^{tm1.1Kge}$) created in the Georgopoulos lab (16). *Chd4* βKO mutant mice were created by breeding *Ins1$^{(Cre/+)}$* mice (B6(Cg)-*Ins1$^{tm1.1(cre)Thor}$*/J) (17) and *Chd4$^{(flox/flox)}$* mice. For experiments requiring cell sorting, *Ins2$^{(GFP/+)}$* reporter mice (18) were bred into the *Chd4* control and βKO line. Physiological studies were conducted on male and female mice with ages of mice being noted for each experiment. Mice were kept in accordance with University of Colorado Institutional Animal Care and Use Committee (IACUC) protocol #00045. Genotyping primers are listed in Supplementary Table 1.

##### *Blood Glucose Readings*

Fasted blood glucose was measured after a 6 hour fast from mouse tail vein. *Ad libitum* blood glucose readings were taken from the mouse tail vein of fed mice. All glucose readings were taken using a Contour 7151H glucose meter and Contour 7097C test strips.

##### *Glucose Tolerance Test*

Mice were fasted for 6 hours and initial blood glucose readings were taken at timepoint 0 from mouse tail vein. Mice were then injected with 2mg per gram of body weight D-glucose (Sigma, Cat. #G8270-1Kg). Additional tail vein blood glucose readings were taken at time 15, 30, 60, 90 and 120 minutes. All glucose readings were taken using a Contour 7151H glucose meter and Contour 7097C test strips. The area under the curve was analyzed using PrismGraphPad 10.4.1.

##### *Mouse Pancreas Tissue Preparation and Immunofluorescence Staining*

Mouse pancreata were dissected at specified ages and fixed in 4% PFA for 4 hours. Following fixation, pancreata were placed in 30% sucrose overnight then embedded in Optimal Cutting Temperature (OCT) Compound (Sakura, Cat. #4583) and frozen at -80°C. Tissue blocks were sectioned using a cryostat (Thermo Microm525) at a slice thickness of 10μm. Three consecutive slices were added to each slide until the entire pancreas was sectioned completely through, ~ 100-120 slides total (300-360 sections per pancreas).

For immunofluorescent staining, every 10th slide, covering the entire pancreas was stained. Prior to staining, slides were thawed for 10 minutes. If antigen retrieval was necessary slides were preheated with Na Citrate (10mM Na Citrate, 0.05% Tween, pH 6.00) and heated for 20 minutes followed by washes using PBS-T (0.1% Tween-20). Slides were then blocked using PBS-T + 2% Normal Donkey Serum (NDS) at room temperature (RT) for 30 minutes. Slides were washed 1x in PBS-T for 5 minutes at RT followed by addition of primary antibodies (Supplementary Table 2). Slides were incubated overnight at 4 degrees in humid chamber. The following day slides were washed 4x in PBS-T for 5 minutes at RT followed by addition of secondary antibodies (1:500 in PBS-T + 2% NDS). Secondary antibodies were incubated on slides for 1-3 hours in humid chamber at RT. Slides were washed 4x in PBS-T for 5 minutes at RT followed by addition of DAPI (1:1000 in PBS-T) and incubated for 15 minutes at RT. Slides were washed 2x in PBS-T for 5 minutes at RT followed by addition of Prolong Gold mounting reagent (Invitrogen P36930), 22x60 cover slips and sealed using clear nail polish (Electron Microscopy Sciences, Cat. #72810).

***Image Analysis and Quantification***

To quantify the P2, 3 week and 6-week timepoints, images were taken on a Leica DM5500B with a Leica DFC 345 FX and Leica DFC425 camera. Filters cubes were A4, L5 and N3. For the P2 timepoint, pictures of entire pancreas sections were taken using the tilescan function using the 10x objective. For the 3-week and 6-week timepoints, pictures of all individual islets were taken using a 20x objective from one of the three pancreas slices of each slide stained. Each islet immunofluorescent image was quantified using ImageJ2 Version 2.14.0/1.54f. Full islet area was quantified as well as positive hormone area by quantifying all area above a set threshold (50 for all channels). Hormone area for P2 was normalized by total pancreas area and hormone area for 3- and 6-week timepoints was normalized by total islet area. For all other immunofluorescent staining, images were taken on a Zeiss LSM 800 with the objective specified and using a 405, 488, 561 and 640 nm laser line.

***Whole Mouse Pancreas Acid Ethanol Insulin Collection***

10-20 mg sections of whole mouse pancreatic tissue were removed from both controls and *Chd4* βKO mutant mice and placed in 1 mL of 0.18M HCl in 70% EtOH (acid ethanol) and sonicated 2 x 30 seconds (Fisher

Scientific Cat. #FS30H) and vortexed for 1 minute. Whole pancreas tissue was incubated at 4°C for 12 hours and stored at -80°C until ELISAs were performed using Mercodia Insulin ELISA (Cat. #10-1247-01) or Mercodia ProInsulin ELISA (Cat. #10-1232-01) per the manufacturer's instructions.

***Generation of Full Mouse Pancreas Slices***

Pancreas tissue slices were prepared as previously described (19). Briefly, animals were injected with 100 mg/kg ketamine and 8 mg/kg xylazine and euthanized via exsanguination prior to inflating the pancreas with low-melting-point 1.25% agarose solution dissolved in extracellular solution (ECS, consisting in 140 mM NaCl, 5 mM KCl, 2 mM $NaHCO_3$, 1 mM $NaH_2PO_4$, 1.2 mM $MgCl_2$, 1.5 mM $CaCl_2$, 3 mM glucose and 10 mM HEPES [pH is adjusted to 7.4 with NaOH]) at 37°C into the common bile duct. Immediately after injection, the agarose infused pancreas was cooled with ice-cold ECS, excised, trimmed into smaller blocks and embedded in 1.25% agarose. Tissues were sliced with a thickness of 200 µm with a vibratome (VF-310-0Z, Precisionary) and collected in ice-cold ECS. Slices were then placed in a 24-well plate containing 1 ml/well of ECS with Soybean Trypsin (Final concentration 0.1 mg/ml) inhibitor to prevent any inadvertent degradation by digestive enzymes. Slices were washed on an orbital shaker at 4°C for 1-hour prior to selection of islet containing slices. Slices were then placed in cell culture inserts coated with a collagen mixture (19) in a 6-well plate with 2 ml of RPMI medium supplemented with 10% FBS, 1% Pen-Strep, and trypsin inhibitor (Final Concentration 0.1 mg/ml), and kept overnight in the Incubator at 37°C and 5% $CO_2$. Slices were then used to perform Glucose Stimulated Insulin Secretion (GSIS) and intracellular calcium imaging experiments.

***Mouse Pancreas Slices Glucose Stimulated Hormone Secretion Assays and Analysis***

Insulin was assayed using static hormone secretion assays. All solutions were prepared in Krebs-Ringer buffer (108.8 mM NaCl, 5 mM $NaHCO_3$, 5.8 mM KCl, 1.2 mM $KH_2PO_4$, 2.5 mM $CaCl_2$, 1.2 mM $MgSO_4$, 10 mM HEPES, 0.1% BSA, pH 7.4) and supplemented with either 2mM, 11mM, or 20 mM Glucose and 20mM, 11 mM, or 2 mM NaCl for osmotic balance. Duplicate sets of 3 or 5 pancreatic slices per condition were incubated in 2 mL of Krebs-Ringer buffer containing 2 mM glucose for 1 hour to establish baseline secretion. For glucose stimulation measurements, each slice set was placed under a sequential 30-minute stimulation condition under one of three conditions: (1) 2 mM glucose, (2) 11 mM glucose, or (3) 20 mM glucose. For GIRK inhibitor measurements, each slice set was placed under a sequential 30-minute stimulation condition under (1) 2 mM glucose, (2) 11 mM or 20 mM glucose, then (3) 20 mM glucose supplemented with either 20 mM KCl or 10 μM VU0468554 (selective GIRK inhibitor; Axon Medchem, Cat. #3593; CAS 1448705-21-2; purity 99%). After each 30-minute stimulation, 500 μL of supernatant was collected for analysis. Samples were stored at -20°C until analysis. Insulin concentrations were measured using a mouse ultrasensitive insulin ELISA kit (Crystal Chem, Cat. #90096) per the manufacturer's instructions.

***Mouse Islet Isolation and Dissociation***

Islets were isolated by the Islet Isolation Core facility. Briefly, mice were anesthetized using a ketamine/xylazine/acepromazine (KXA) mixture and euthanized by exsanguination. The pancreatic duct was clamped off and a 26-gauge needle with a collagenase mixture was injected into the pancreas. The pancreas was dissected out of the mouse and islets were isolated by a density gradient purification step followed by hand picking.

After isolation, islets were collected and placed in a 15mL tube and allowed to settle for 20 minutes. Islets were washed with pre-warmed PBS and spun at 300 x g for 3 minutes at RT. PBS was aspirated off and islets were resuspended in 2 mL of Accutase (Stemcell Tech, Cat. #07920) and placed in a 37°C water bath. Islets were incubated for 30 minutes total, resuspending islets by pipetting every 5 minutes. Accutase dissociation was stopped by adding RPMI1640 (Gibco, Cat. #11875-093) + 10% FBS. Dissociated islet cells were filtered through a 40μm cell strainer (Greiner, Cat. #542040) into a 50mL conical tube. Cells were spun at 300 x g for 3 minutes at RT. Supernatant was carefully aspirated off without disturbing cell pellet and cells were washed in 10mL FACS buffer (PBS + 1mM EDTA + 0.2% BSA). Cells were spun at 300 x g for 3 minutes at RT, supernatant was aspirated off and cells were resuspended in 1mL of FACS buffer for sorting. Propidium Iodide (PI, Thermo Cat. #P3566) was added as a live/dead marker at 1:1000.

***Mouse Islet β Cell Fluorescent Activated Cell Sorting (FACS)***

β cells were sorted from dissociated 4 week mouse islets using an *Ins2*$^{(GFP/+)}$ reporter (18) on a Bio-rad S3e cell sorter. Gates were set to initially read FSC and SSC area to sort out debris and cell doublets. Cells were then gated using a PI negative to separate out living vs dead cells and lastly cells were gated using a GFP positive gate to sort and collect all β cells (Supplementary Fig. 5). Cells were collected in 350 µL of RLT Lysis buffer + βME (Qiagen RNeasy Micro Kit, Cat. #74004), prior to RNA extraction.

***RNA Extraction and RNA Sequencing***

RNA extraction was performed using the RNeasy Micro Kit (Qiagen, Cat. 74004) protocol. 2-Mercaptoethanol (βME) was added to RLT lysis buffer to help inhibit RNases. RNA quality was assessed using an Agilent 4200 TapeStation System and RNA samples with RNA Integrity Numbers (RIN) scores of 8 or higher were used for sequencing. RNA samples were sequenced at the Genomics Core facility on the University of Colorado Anschutz Medical Campus. Paired end libraries were made by the core from low input ribo-depleted RNA using SMARTer Stranded Total RNA-Seq Kit v2 – Pico Input Mammalian (Takara Cat.# 634411). All libraries were run on a NovaSEQ X machine at a sequencing depth of 50 million read pairs / 100 million total reads per sample. Fastq files were quality checked using FastQC (v0.12.1) (20) and adapters were trimmed using Cutadapt (v4.8) (21). The reads were mapped to the mm10 genome using STAR (v2.7.11a) (22). Counts tables were made using featureCounts (v2.0.6) (23) and used for differential gene expression analysis. Differential genes were assessed using DESeq2 (v1.46.0) (24) and significant genes were established using a padj value of $\leq$ 0.05 and a $Log_2$ Fold Change value of > 0.5 or < 0.5.

***Reanalysis of ATAC-seq Dataset***

The reanalysis of the published ATAC-seq data (25) was conducted by downloading Fastq files from the GEO database under reference number GSE217444. Files were processed using our bulk ATAC-seq snakemake pipeline. Briefly, Fastq files were quality checked using FastQC (v0.12.1) (20) and adapters were trimmed using Cutadapt (v4.8) (21). The reads were aligned to the mm10 genome using Bowtie2 (v2.5.3) (26) and mitochondrial reads were removed using Samtools (v1.20) (27). The ATACseqQC package (v1.28) (28) was used to generate final quality metric plots, and FRiP values were tabulated using featureCounts (v2.0.6) (23). We used MACS3 (v3.0.2) (29) with a q-value cutoff of 0.01 to call peaks within the dataset, then utilized DiffBind (v3.14) (30) with DESeq2 (v1.46.0) (24) and a padj value < 0.05 to identify differentially accessible peaks between the CHD4 Control and CHD4 βKO ATAC-seqs.

***CHD4 ChIP-qPCR***

20 million MIN6 cells were collected for use in the SimpleChIP Enzymatic Chromatin IP Kit (Magnetic Beads, CST, Cat. #9003) per the manufacturer's instructions. Briefly, cells were crosslinked with 1% formaldehyde for 10 minutes and crosslinking was quenched with glycine. Cells were lysed and chromatin was fragmented using a Diagenode Bioruptor Pico sonicator and Micrococcal Nuclease (CST, Cat. # 10011) to fragment chromatin to approximately 150-900bp. 2% of samples was removed to be used as input before pulling down CHD4. 30 µg of fragmented chromatin DNA was used for IP with 10 µL CHD4 antibody (Rabbit, CST D4B7, 12011S) or 2µL IgG (Rabbit, CST, Cat. #2729). Samples were next incubated overnight with rotation at 4°C and 30 µL of ChIP-Grade Protein G Magnetic Beads (CST, Cat. #9006) was added, followed by incubating samples again for 2 hours with rotation at 4°C. Chromatin was eluted from beads and cross-links were reversed using NaCl, RNase A and Proteinase K and DNA was purified from samples using DNA spin columns.

For qPCR, 3 replicates of 5 ng of CHD4 IP or IgG IP samples were used with iTaq Universal SYBR Green Supermix (Bio-Rad, Cat. #1725120) and primers (Supplementary Table 1). A Bio-Rad CFX96 system was used to run qPCR plate. Fold enrichment over IgG was used to quantify CHD4 binding over IgG control.

***Mouse Pancreas Slices Calcium Signaling and Analysis***

Slices were incubated with 4 µM Fluo-4 AM (Thermo Fisher Scientific Cat. #F14201) in BMHH buffer containing 125 mM NaCl, 5.7 mM KCl, 2.5 mM $CaCl_2$, 1.2 mM $MgCl_2$, 10 mM HEPES, 0.1% bovine serum albumin (BSA), 2 mM glucose, and 1mg/ml Trypsin inhibitor at room temperature for 30 min in the dark. Slices were transferred to MatTek glass-bottom imaging dishes (MatTek, Cat. #P35G-1.5-14-C) with fresh BMHH with 2 mM glucose prior to imaging. An anchor was placed to hold the slice in place. Slices were imaged on a Zeiss laser scanning microscope (LSM) 800 with a 40x objective using a 488 nm laser. Slices were imaged at 37°C, where images were acquired every second for 2.5–5 min at 2mM glucose. Following imaging, medium was changed to 11 mM glucose and images were acquired for another 15-30 min. Fiji was used to extrapolate the calcium time courses. The time courses were normalized for intensity and normalized for movement if needed. The area under the curve was analyzed using PrismGraphPad 10.4.1.

***Statistical Analysis***

Unless otherwise stated, error bars are shown as mean +/- standard deviation (SD) and Shapiro-Wilk test was used to test normality. If passed, unpaired two-tailed student's t-test was used to determine significant differences, otherwise, Mann-Whitney U test was used. P-value or padj of $\leq 0.05$ was used to determine statistical significance. If no statistics are shown on graphs, results were not significant.

***Data and materials availability***

The datasets and computer code produced in this study are available in the following databases:

- RNA-sequencing data: Gene Expression Omnibus GSE299270 (https://www.ncbi.nlm.nih.gov/geo/query/acc.cgi?acc=GSE299270)
- ATAC-sequencing data: Gene Expression Omnibus GSE217444 (https://www.ncbi.nlm.nih.gov/geo/query/acc.cgi?acc=GSE217444).
- Code for the analysis of the RNA sequencing data is available on Github (https://github.com/CUAnschutzBDC/snakemake_pipelines/tree/6ea22de25f49706bb268502bbd894a4a4b7cdb55/RNA_seq)
- Code for the analysis of the ATAC sequencing data is available on Github (https://github.com/CUAnschutzBDC/snakemake_pipelines/tree/6ea22de25f49706bb268502bbd894a4a4b7cdb55/ATAC_seq)
- Docker containers for the RNA-seq are available on docker hub: (https://hub.docker.com/layers/kwellswrasman/rnaseq_general/v1/images/sha256-2536b14130bba19b70f28d23807f127854cd646bbc4bd4028407eeb35ffea45f ), (https://hub.docker.com/layers/kwellswrasman/rnaseq_r/v1/images/sha256-36d320d7de293da2a84ff35ab131ef6ebec42ab53934aacc9d4f341f7da33988).
- Docker containers for the ATAC-seq are available on docker hub: (https://hub.docker.com/layers/kwellswrasman/fastq_screen/v1/images/sha256-cf697c13d436f251dc70048adfee58438576698c03e398455c394c2d16ffac59), (https://hub.docker.com/layers/kwellswrasman/general_chip/v2/images/sha256-447cd6c6ab57006b75b285f851443b9b9d0bc06c8d882bb1a1cb3387d61cc499), (https://hub.docker.com/layers/kwellswrasman/atac_chip_r/v1/images/sha256-e499fc2bcde315a720a442447decb95c3a4fa084466bdf816e164f2e40f01971), (https://hub.docker.com/repository/docker/kwellswrasman/picard/tags/v1/sha256-29984d83a5ada3c1799684c36bfb4a1dce6912a105baa3b87828b3ef939ee7de).

### Supplemental Figures

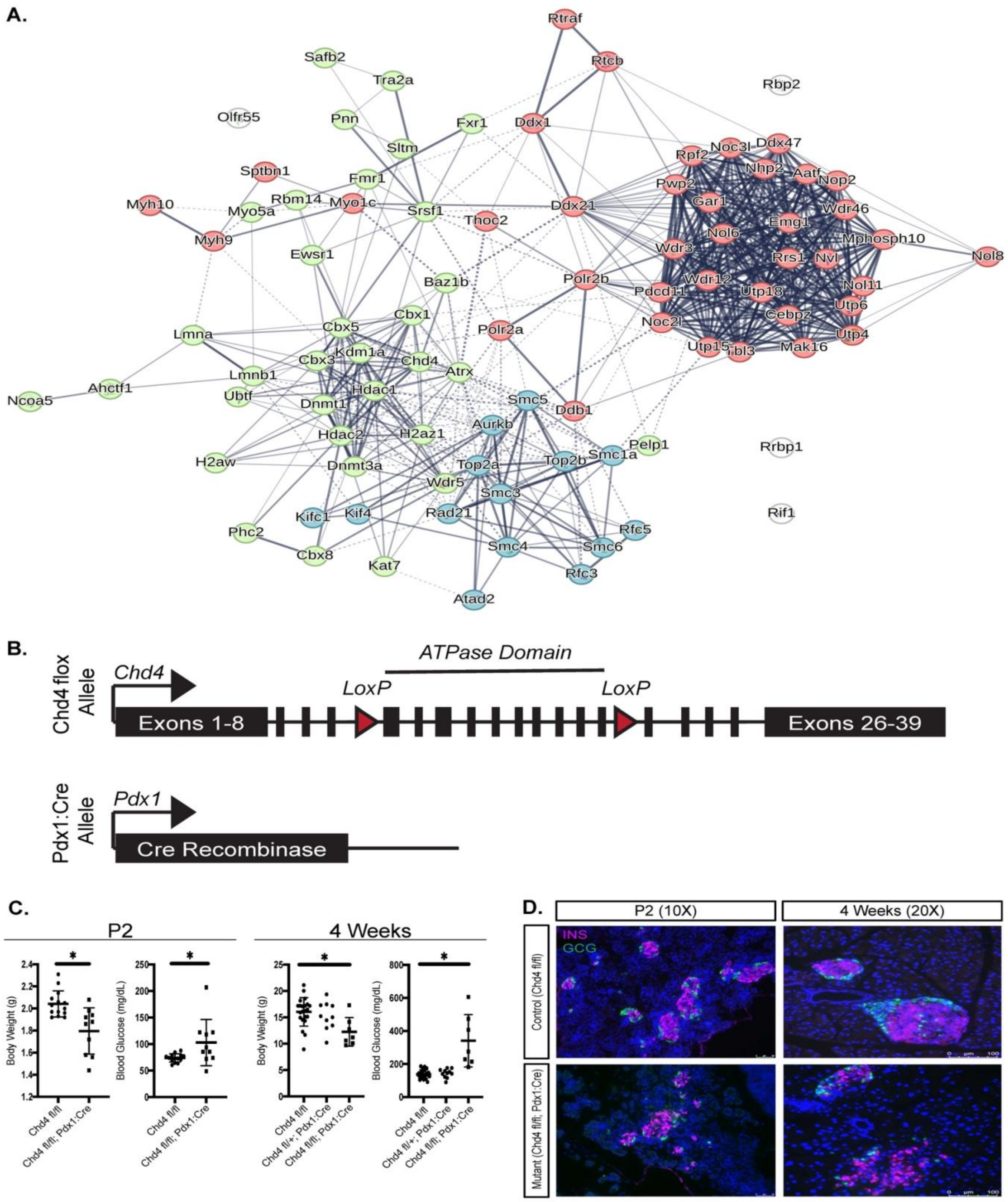

**Supplementary Figure 1.**

**Deletion of *Chd4* from the developing pancreas causes hyperglycemia and body weight defects.** (A) String database clustering identified 112 overlapping proteins from NKX2.2 IP-MS and CHD4 IP-MS that grouped into 3 k-means clusters (18). (B) Schematic of Pancreas and Duodenum *Chd4* KO allele. Flanking Lox P (flox) sites are located upstream of exon 12 and downstream of exon 21, excising the ATPase domain. (C) Measurements of body weight and *ad libitum* blood glucose for P2 control ($Chd4^{fl/fl}$, n =14) and mutant ($Chd4^{fl/fl}$; $Pdx1^{Cre/+}$, n=10) mice (two-tailed Student's *t*-test). Measurements of body weight and *Ad libitum* blood glucose for 4-week control ($Chd4^{fl/fl}$, n=22), Het ($Chd4^{fl/+}$; $Pdx1^{Cre/+}$, n=10) and mutant ($Chd4^{fl/fl}$; $Pdx1^{Cre/+}$, n=7) mice (one-way ANOVA test). (D) Representative immunofluorescent images for P2 and 4-week control and mutant mice showing DAPI (blue), Insulin (INS, magenta), and Glucagon (GCG, green). P2 IF images are at 10x magnification with Scale bars: 50 µm. 4-week IF images are at 20x magnification with Scale bars: 100 µm. *$P \leq 0.05$.

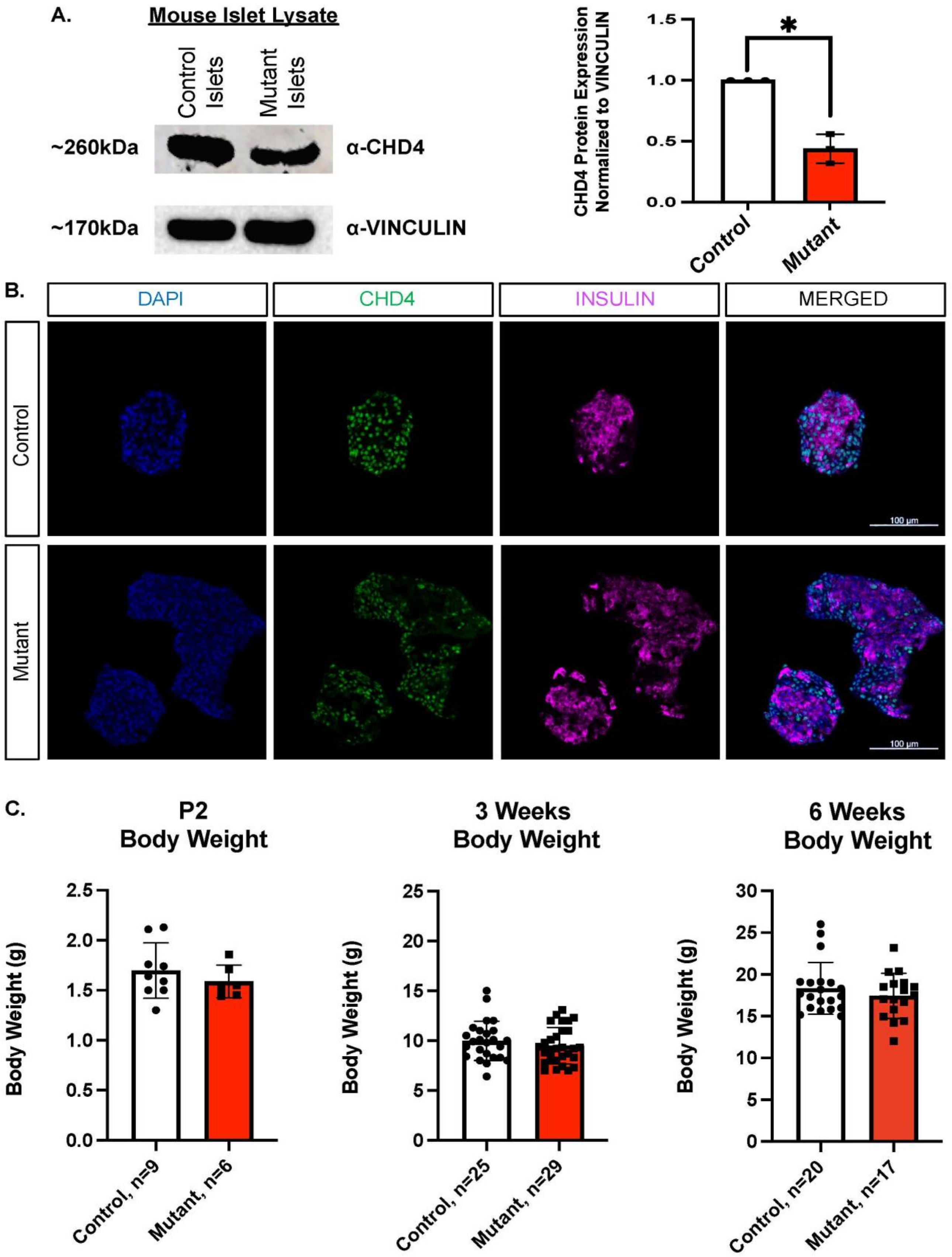
A.
Mouse Islet Lysate
Control Islets
Mutant Islets
~260kDa
α-CHD4
~170kDa
α-VINCULIN
CHD4 Protein Expression Normalized to VINCULIN
1.5
1.0
0.5
0.0
*
Control
Mutant
B.
DAPI
CHD4
INSULIN
MERGED
Control
Mutant
100 μm
100 μm
C.
P2 Body Weight
Body Weight (g)
2.5
2.0
1.5
1.0
0.5
0.0
Control, n=9
Mutant, n=6
3 Weeks Body Weight
Body Weight (g)
25
20
15
10
5
0
Control, n=25
Mutant, n=29
6 Weeks Body Weight
Body Weight (g)
30
25
20
15
10
5
0
Control, n=20
Mutant, n=17

**Supplementary Figure 2.**

***Chd4* is efficiently deleted from mouse islet β cells without a disruption in body weight.** (A) Western blot analysis of CHD4 expression in mouse islet lysate. Quantification of CHD4 expression normalized to Vinculin between control and mutant lysates (two-tailed Student's *t*-test). (B) Representative immunofluorescent images (20x magnification) for 3-week control and mutant mice showing *Chd4* KO. DAPI (blue), Insulin (magenta), and CHD4 (green). Scale bars: 100 µm. (C) Measurements of body weight for P2, control (n=9) and mutant (n=6) mice (two-tailed Student's *t*-test). Measurements of body weight for 3-week, control (n =25) and mutant (n=29) mice (two-tailed Student's *t*-test). Measurements of body weight for 6-week, control (n =20) and mutant (n=17) mice (Mann Whitney U test). *$P \leq 0.05$.

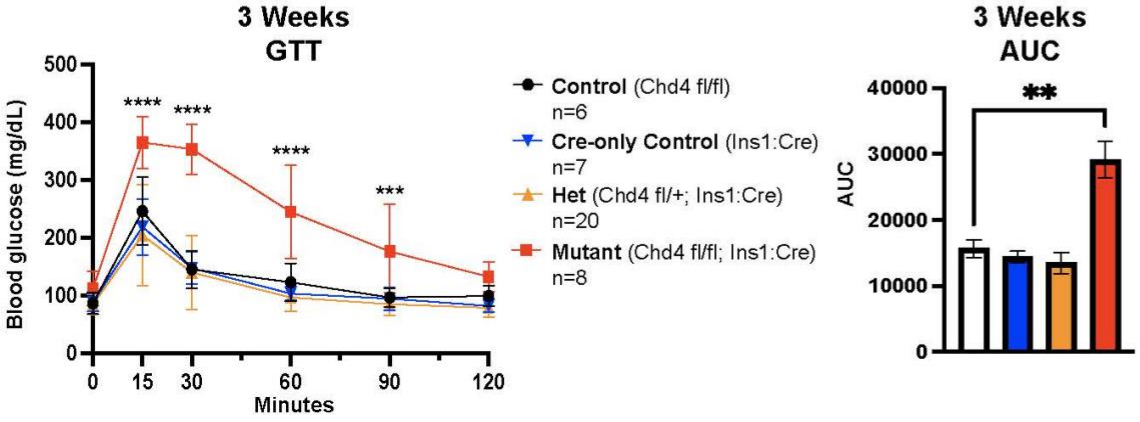
3 Weeks
GTT
Blood glucose (mg/dL)
500
400
300
200
100
0
****
****
****
***
0
15
30
60
90
120
Minutes
Control (Chd4 fl/fl)
n=6
Cre-only Control (Ins1:Cre)
n=7
Het (Chd4 fl/+; Ins1:Cre)
n=20
Mutant (Chd4 fl/fl; Ins1:Cre)
n=8
3 Weeks
AUC
AUC
40000
30000
20000
10000
0
**

**Supplementary Figure 3.**

**All control genotypes have similar fasted blood glucose and glucose tolerance levels.**

3-week combined sex Glucose Tolerance Test (GTT) of control (*Chd4*$^{fl/fl}$ : black line, n=6), Cre-only control (*Ins1*$^{Cre/+}$: blue line, n=7), het (*Chd4*$^{fl/+}$*; Ins1*$^{Cre/+}$: orange line, n=20), and mutant (*Chd4*$^{fl/fl}$*; Ins1*$^{Cre/+}$: red line, n=8) mice (multiple comparison with 2 way ANOVA). Area under the curve (AUC) measurement for respective 3-week GTT. All GTT points normalized to 0 minutes time point prior to AUC calculation (one-way ANOVA). $^{**}P \leq 0.01$, $^{***}P \leq 0.001$, $^{****}P \leq 0.0001$.

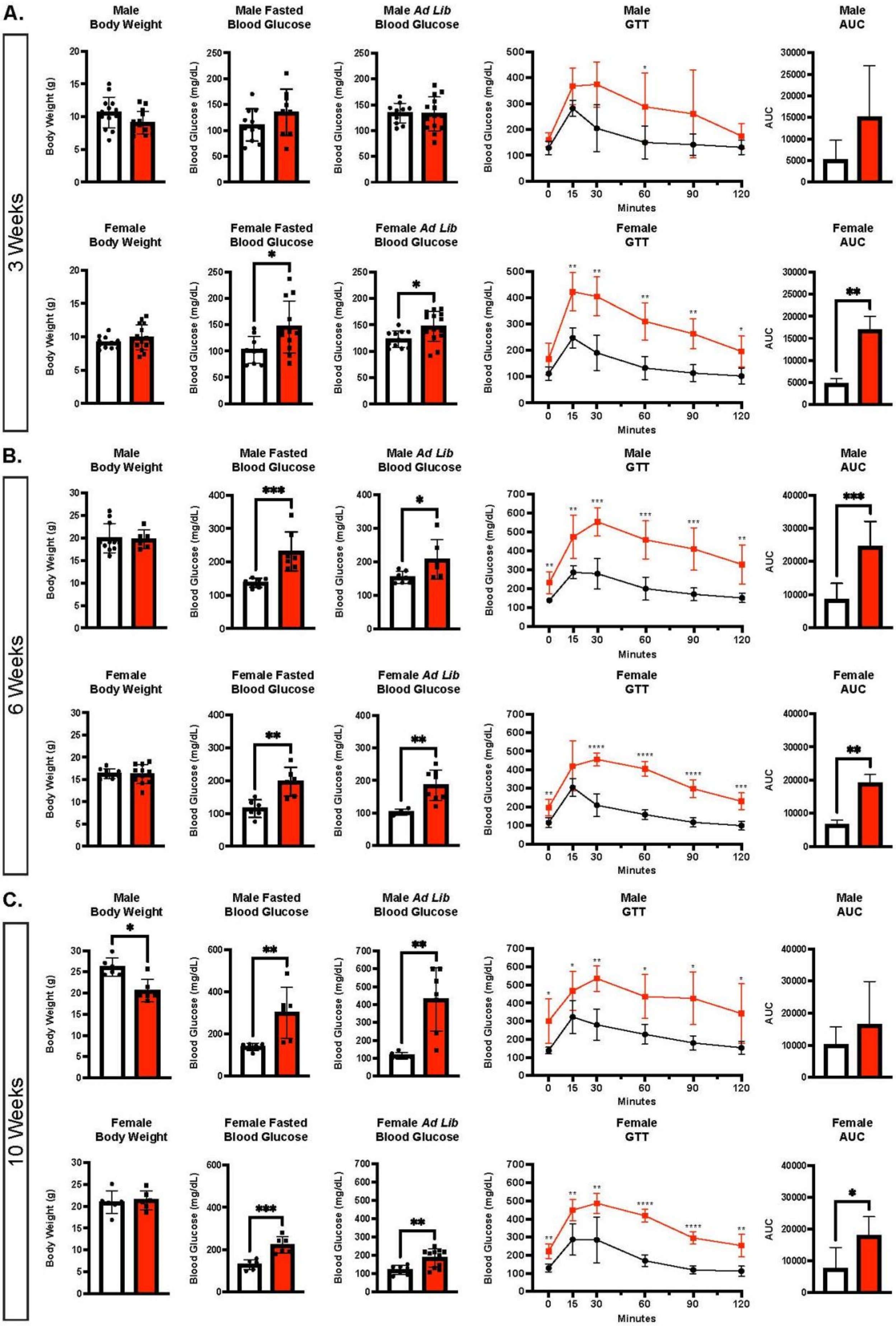
A.
3 Weeks
Male Body Weight
Male Fasted Blood Glucose
Male Ad Lib Blood Glucose
Male GTT
Male AUC
Female Body Weight
Female Fasted Blood Glucose
Female Ad Lib Blood Glucose
Female GTT
Female AUC
B.
6 Weeks
Male Body Weight
Male Fasted Blood Glucose
Male Ad Lib Blood Glucose
Male GTT
Male AUC
Female Body Weight
Female Fasted Blood Glucose
Female Ad Lib Blood Glucose
Female GTT
Female AUC
C.
10 Weeks
Male Body Weight
Male Fasted Blood Glucose
Male Ad Lib Blood Glucose
Male GTT
Male AUC
Female Body Weight
Female Fasted Blood Glucose
Female Ad Lib Blood Glucose
Female GTT
Female AUC
Body Weight (g)
Blood Glucose (mg/dL)
AUC
Minutes

**Supplementary Figure 4.**

**Both male and female *Chd4* βKO mice show a worsening diabetic phenotype.**

Unless otherwise noted, mouse data consists of control (*Chd4*$^{fl/fl}$ : white Bars) and mutant/*Chd4* βKO (*Chd4*$^{fl/fl}$; *Ins1*$^{Cre/+}$: red Bars) genotypes. (A) Measurements of 3-week male control (n =14) and mutant (n=14*)* body weight. 3-week female control (n =11) and mutant (n=15*)* body weight. 3-week male control (n =11) and mutant (n=9*)* fasted blood glucose. 3-week female control (n =9) and mutant (n=12*)* fasted blood glucose. 3-week male control (n =11) and mutant (n=14*) Ad libitum (Ad Lib)* blood glucose. 3-week female control (n =11) and mutant (n=14*) Ad Lib* blood glucose. 3-week male Glucose Tolerance Test (GTT) of control (black line, n=6) and mutant (red line, n=6) mice. 3-week female GTT of control (black line, n=6) and mutant (red line, n=6) mice. Area under the curve (AUC) measurement for respective 3-week GTTs. All points normalized to 0 minutes time point prior to AUC calculation. (B) Measurements of 6-week male control (n =11) and mutant (n=6*)* body weight. 6-week female control (n =9) and mutant (n=11*)* body weight. 6-week male control (n =7) and mutant (n=7*)* fasted blood glucose. 6-week female control (n =6) and mutant (n=6*)* fasted blood glucose. 6-week male control (n =7) and mutant (n=6*) Ad Lib* blood glucose. 6-week female control (n =4) and mutant (n=8*) Ad Lib* blood glucose. 6-week male GTT of Control (black line, n=7) and mutant (red line, n=7) mice. 6-week female GTT of control (black line, n=6) and mutant (red line, n=6) mice. Area under the curve (AUC) measurement for respective 6-week GTTs. All points normalized to 0 minutes time point prior to AUC calculation. All points normalized to 0 minutes time point prior to AUC calculation. (C) Measurements of 10-week male control (n =6) and mutant (n=6*)* body weight. 10-week female control (n =6) and mutant (n=6*)* body weight. 10-week male control (n =6) and mutant (n=6*)* fasted blood glucose. 10-week female control (n =6) and mutant (n=6*)* fasted blood glucose. 10-week male control (n =6) and mutant (n=7*) Ad Lib* blood glucose. 10-week female control (n =7) and mutant (n=11*) Ad Lib* blood glucose. 10-week male GTT of control (black line, n=6) and mutant (red line, n=6) mice. 6-week female GTT of control (black line, n=6) and mutant (red line, n=6) mice. Area under the curve (AUC) measurement for respective 10-week GTTs. All points normalized to 0 minutes time point prior to AUC calculation. For all panels in figure, *$P \leq 0.05$, **$P \leq 0.01$, ***$P \leq 0.001$, ****$P \leq 0.0001$. For GTT figures, multiple two-tailed Student's *t*-tests were used. For all other figures, two-tailed Student's *t*-test except Male 6-week body weight, Male 6-week fasted blood glucose, Male 10-week body weight, and Male 10-week fasted blood glucose where Mann Whitney U tests were used.

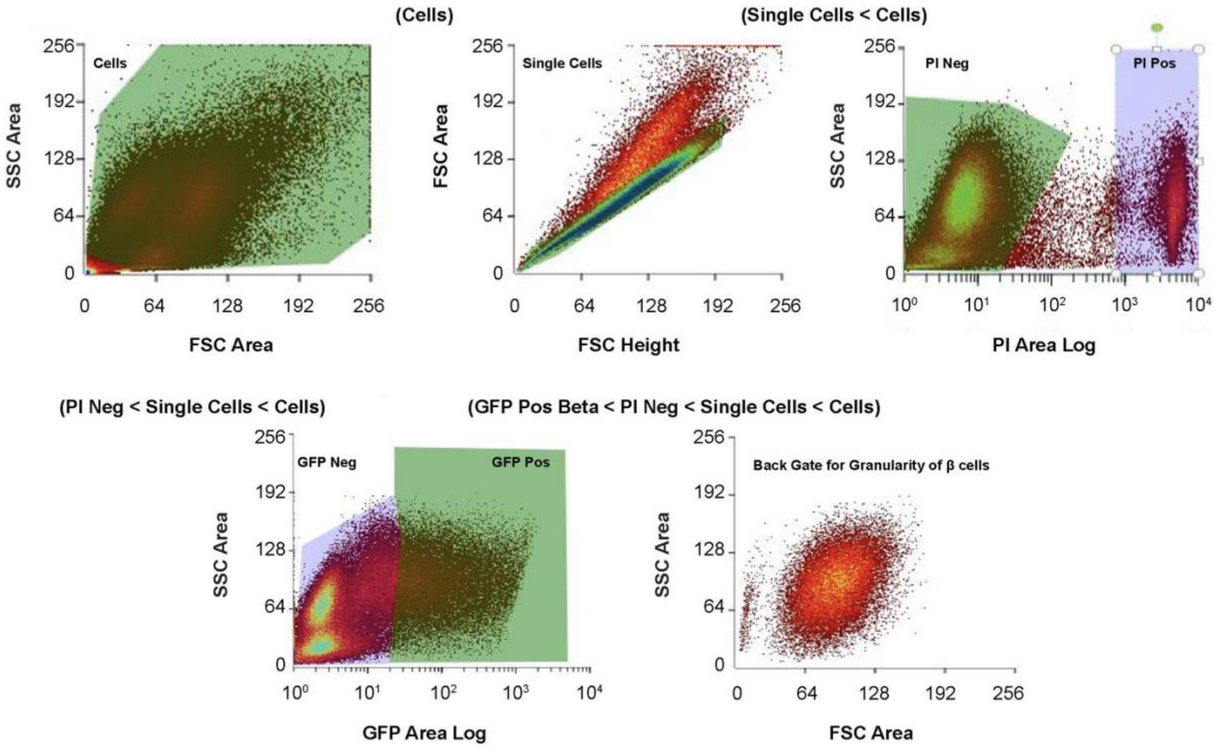

(Cells)
(Single Cells < Cells)
Cells
Single Cells
PI Neg
PI Pos
SSC Area
FSC Area
FSC Height
PI Area Log
(PI Neg < Single Cells < Cells)
(GFP Pos Beta < PI Neg < Single Cells < Cells)
GFP Neg
GFP Pos
Back Gate for Granularity of β cells
GFP Area Log

**Supplementary Figure 5.**

**Gating used to sort β cells using GFP.** β cells were sorted using an *Ins2*$^{(GFP/+)}$ reporter allele, bred into control and mutant mice. Cells were initially sorted using FSC area and SSC area to remove debris and doublet cells. Cells were next sorted using propidium iodide (PI) as a live/dead marker, gating on live cells. Cells were next sorted using a GFP+ area. Lastly, cells were back gated to FSC area and SSC area to check granularity of cells to make sure they replicate β cell granularity.

A.

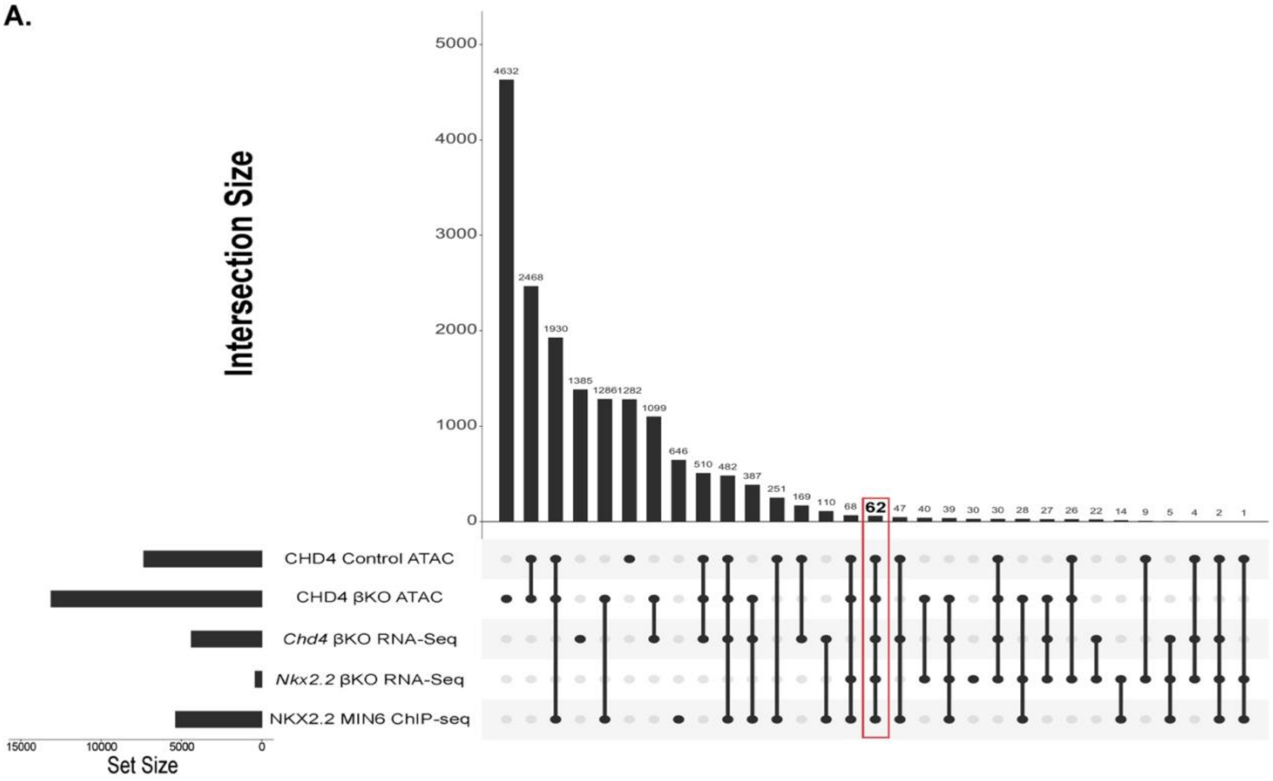


B.

| Gene | Base_mean_Chd4KO | Log2FC_Chd4KO | padj_Chd4KO | Base_mean_Nkx2.2KO | Log2FC_Nkx2.2KO | padj_Nkx2.2KO |
|---|---|---|---|---|---|---|
| Kcnj5 | 1872.765351 | 6.498529061 | 5.9353E-302 | 660.9224084 | 7.575281622 | 1.83396E-13 |
| Ambp | 4210.534507 | 3.450108275 | 1.8596E-265 | 550.0779496 | 1.110022203 | 3.73349E-07 |
| Ucn3 | 4696.286527 | -2.908218492 | 9.9507E-112 | 7335.524557 | -1.177566172 | 1.8113E-09 |
| Clmn | 644.4948888 | 3.17352536 | 1.7819E-106 | 256.7113024 | 1.308047612 | 1.88532E-08 |
| Cacna2d2 | 424.1499709 | 3.974684735 | 6.04529E-83 | 194.4222888 | 1.302511869 | 0.000130809 |
| Ttc28 | 5362.049956 | -1.270957972 | 1.56675E-59 | 2390.463826 | -0.581562641 | 0.002084447 |
| Slc2a2 | 5907.137065 | -1.748390054 | 3.77372E-54 | 18372.53568 | -1.101644286 | 1.9265E-09 |
| Myo1b | 3310.314887 | 1.698182873 | 3.70976E-50 | 1357.822974 | 0.711423037 | 0.015144027 |
| Lrrn1 | 4372.183785 | -1.219832147 | 1.95409E-46 | 4007.22846 | -0.681397832 | 0.000779483 |
| Itgav | 9796.630177 | -1.044232345 | 1.99529E-46 | 2997.364376 | -0.657909589 | 0.003831996 |
| Fmnl2 | 4232.658315 | -1.61625206 | 6.56776E-45 | 1716.898649 | -0.757008495 | 3.20847E-06 |
| Cadm1 | 2081.018928 | 2.02489776 | 2.02451E-41 | 869.8251045 | 0.994195538 | 8.4141E-05 |
| Basp1 | 320.8731209 | 2.798729676 | 3.44469E-36 | 247.5484394 | 1.208907829 | 0.003040651 |
| Plekha1 | 4127.149357 | 0.946387644 | 5.79587E-31 | 4015.011652 | 0.866156153 | 1.90179E-05 |
| Cdh8 | 1169.686207 | -1.664937007 | 2.1677E-27 | 1440.438854 | -0.65005154 | 0.000674251 |
| Gprc5b | 166.953635 | 2.708880349 | 3.79327E-26 | 327.3267084 | 0.757813189 | 0.000698191 |
| Slc30a8 | 6952.746943 | -0.934357201 | 3.6468E-22 | 18082.30726 | -0.717740244 | 3.14437E-12 |
| Anks1b | 100.2501844 | 3.985308199 | 8.94996E-20 | 46.21137644 | 1.858267313 | 0.001738487 |
| Pcx | 6132.71992 | -0.884936928 | 4.59166E-19 | 2712.260804 | -0.873499811 | 5.106E-09 |
| Tmem150c | 1017.337418 | -1.088107973 | 1.10533E-17 | 580.732341 | -0.740654084 | 0.010043715 |
| G6pc2 | 26605.61363 | -0.980184341 | 2.20465E-17 | 50701.50734 | -0.674291635 | 0.015528308 |
| Sdk2 | 28092.81237 | -0.744523214 | 7.66544E-17 | 2658.65845 | -0.601551283 | 0.000832627 |
| Phactr1 | 16141.74687 | -0.676308498 | 2.0065E-16 | 4096.827462 | -0.800810132 | 0.000155413 |
| Lifr | 3730.40416 | -0.729827446 | 4.48637E-16 | 2253.504361 | -0.771784173 | 7.08736E-09 |
| Arrdc4 | 882.5439532 | 0.922058315 | 2.18312E-15 | 646.4945731 | 0.934672123 | 0.000105988 |
| Idh2 | 4116.009818 | -0.759237967 | 5.54155E-15 | 2559.117653 | -0.548088162 | 0.004881916 |
| Sez6l | 18436.21582 | -1.044762333 | 2.0929E-14 | 3378.853649 | -0.566446919 | 2.8024E-05 |
| Dennd4c | 6742.534448 | -0.678021864 | 1.08508E-13 | 4131.830686 | -0.624453046 | 0.00073531 |
| Reln | 149.9877752 | -2.386250722 | 3.57947E-13 | 73.13701324 | -1.79794385 | 0.00341967 |
| Car10 | 1470.258567 | -0.96589574 | 4.06706E-13 | 1181.594146 | -0.74878094 | 0.022279322 |
| Adarb1 | 1689.717807 | -0.754214144 | 1.50906E-10 | 937.6592775 | -0.76102876 | 0.001762971 |
| Nrxn1 | 3270.244063 | -0.558664902 | 2.53481E-08 | 902.6119755 | -0.668160059 | 0.01261327 |
| Rhobtb1 | 2859.317037 | -0.616292928 | 2.2099E-07 | 2551.564103 | -0.620162936 | 1.63238E-06 |
| Tle4 | 405.4050288 | 0.829933068 | 6.68935E-07 | 366.8917554 | 1.336221352 | 1.60642E-08 |
| Dscam | 98.68047946 | 1.467706757 | 7.36257E-06 | 173.15051 | 0.923639207 | 0.011588293 |
| Kctd8 | 329.9150575 | 0.749341337 | 3.68597E-05 | 147.9072707 | 0.879626547 | 0.003024339 |
| Ovol2 | 220.3114851 | 0.904531449 | 8.73773E-05 | 272.1818376 | 1.464197519 | 4.4292E-08 |
| Npy | 2980.832948 | 2.573780855 | 0.001523512 | 1084.894626 | 3.162309684 | 6.64648E-23 |
| Irak3 | 104.7186658 | 0.939893311 | 0.005134821 | 81.53852898 | 1.062084019 | 0.035094661 |
| Tcim | 195.6825654 | 0.772921917 | 0.009124929 | 505.9970286 | 0.695144998 | 0.022289545 |

**Supplementary Figure 6.**

**Discovering potential direct targets of CHD4 using overlapping datasets .** (A) UpSet plot showing all the genes that overlap in various combinations throughout the five datasets, *Chd4* control and βKO ATAC-seq, *Chd4* and *Nkx2.2* βKO RNA-seq and *Nkx2.2* ChIP-seq datasets. 62 genes overlapped in all five datasets. (B) List of 40 genes that were dysregulated in the same direction between the *Chd4* and *Nkx2.2* βKO RNA-seqs, with Base Mean, Log2FC and padj values of each gene. List is ordered by smallest to largest padj value from *Chd4* βKO RNAseq.
